# Genomic indicators of gene function: A systematic assessment of the human genome

**DOI:** 10.64898/2026.04.08.717348

**Authors:** Helena B. Cooper, Karla E. Rojas López, Daniela Schiavinato, Michael A. Black, Paul P. Gardner

## Abstract

Proteins and non-coding RNAs are functional products of the genome that are central for crucial cellular processes. With recent technological advances, researchers can sequence genomes in the thousands and probe numerous genomic activities of many species and conditions. Such studies have identified thousands of potential proteins, RNAs and associated activities. However there are conflicting interpretations of the results and therefore which regions of the genome are “functional”. Here we investigate the relative strengths of associations between coding and non-coding gene functionality and genomic features, by comparing reliably annotated functional genes to non-genic regions of the genome. We find that the strongest and most consistent association between functional genes and genomic features are transcriptional activity **and** evolutionary conservation. We also evaluated sequence-based statistics, genomic repeats, epigenetic and population variation data. Other features strongly associated with function include histone marks, chromatin accessibility, genomic copy-number, and sequence alignment statistics such as coding potential and covariation. We also identify potential issues with SNP annotations in short non-coding RNAs, as some highly conserved ncRNAs have significantly higher than expected SNP densities. Our results demonstrate the importance of evolutionary conservation and transcription activity for indicating protein-coding and non-coding gene function. Both should be taken into consideration when differentiating between functional sequences and biological or experimental noise.

## Introduction

A substantial portion of the human genome – particularly intergenic and intronic regions – is composed of repeat-derived, non-protein-coding sequences that remain poorly understood. These regions often evolve at neutral rates and appear to contribute little to organismal fitness. As a result, they have historically been labelled as “junk DNA” (Germain et al. 2014; Doolittle 2013; Doolittle and Brunet 2017; Rands et al. 2014). The term serves not only as a descriptor but also as a null hypothesis in genomics, where function must be demonstrated, not assumed (Koonin 2016).

This perspective stands in contrast to claims from projects such as ENCODE, which argue that widespread biochemical activity, particularly transcription, is indicative of function (ENCODE Project Consortium 2012; ENCODE Project Consortium et al. 2020; Dinger et al. 2009; Mattick et al. 2010). However, critics argue that biochemical activity alone is insufficient, as it can arise from biological noise, leaky transcription, or experimental artifacts. According to the “selected effect” definition of function, activity must also be associated with fitness effects or evolutionary constraint (Doolittle et al. 2014; Doolittle and Brunet 2017; Eddy 2012; Palazzo and Gregory 2014; Germain et al. 2014).

Indeed, numerous studies (Christmas et al. 2023; Haerty and Ponting 2014) have shown that much of the genome is evolutionarily indistinguishable from neutral sites such as ancestral repeats or four-fold degenerate sites, providing empirical support for this more conservative view. These findings reinforce the need to combine evolutionary and biochemical evidence to confidently define functional regions.

Defining functional elements in the human genome remains challenging (Tsai et al. 2017; Kellis et al. 2014). Many observed biochemical activities, such as transcription, may simply reflect biological noise or experimental artifacts (Raser and O’Shea 2005; Eddy 2012; Doolittle 2013; Graur 2017). Random transcription can arise from non-specific DNA-binding proteins, indiscriminate promoters, or transcribed transposons, and random sequences are increasingly shown to yield bioactive RNAs or peptides (Raser and O’Shea 2005; Struhl 2007); (Frumkin and Laub 2023; Hlouchova 2023; Fajardo 2021; Dube et al. 1993; Luthra et al. 2024). Ongoing efforts are working to establish baseline biochemical activity in the absence of selection (Eddy 2024).

Biochemical activity alone does not resolve the C-value paradox, where genome size varies widely without corresponding changes in phenotypic complexity. For example, closely related vizcacha rats differ twofold in genome size (Evans et al. 2017). This disparity is largely explained by the accumulation of selfish, repeat-derived sequences, particularly transposable elements (TEs), which evolve predominantly under neutral drift (Osmanski et al. 2023); (Arkhipova 2018). While some TEs have been co-opted for host function, generalising from a few well-characterised cases to widespread functionality risks confirmation bias and circular reasoning (Bourque et al. 2018; Hayward and Gilbert 2022; Doolittle and Brunet 2017; Lowe and Haussler 2012).

We hypothesise that functional coding and non-coding genes can be distinguished from the genomic background, which is largely composed of decayed or inactive repeat elements (Klapproth et al. 2021; Frankish et al. 2019; Bourque et al. 2018). To test this, we systematically evaluate a broad set of genomic features previously linked to functionality, using consistent and controlled statistical comparisons. Our analysis highlights the most informative signals for defining function and helps clarify the criteria needed to robustly distinguish genic regions from non-functional DNA.

## Results

Our study assesses the statistical relevance of six classes of genomic features for determining gene functionality. We include sequence conservation, transcription levels, epigenetic signatures, genomic repeat associations, protein and RNA-specific scores, and population variation (Figure 1). We compare these features across three main categories of genes – protein-coding (mRNA), short non-coding RNA (sncRNA), and long non-coding RNA (lncRNA) – against length-matched non-genic control regions. The selected genomic features have independently been associated with function in previous analyses and were selected from a range of statistics and scores that met the following inclusion criteria: readily accessible for human genome version GRCh38/hg38, are genome-wide, unbiased to known genes and not redundant (Supplementary Table S1 and S2 for further details of included and excluded features; see also spearman correlation matrices Figures S1-3). Here we present each analysis and discuss the biological relevance to the definition of functionality.

**Figure 1.**
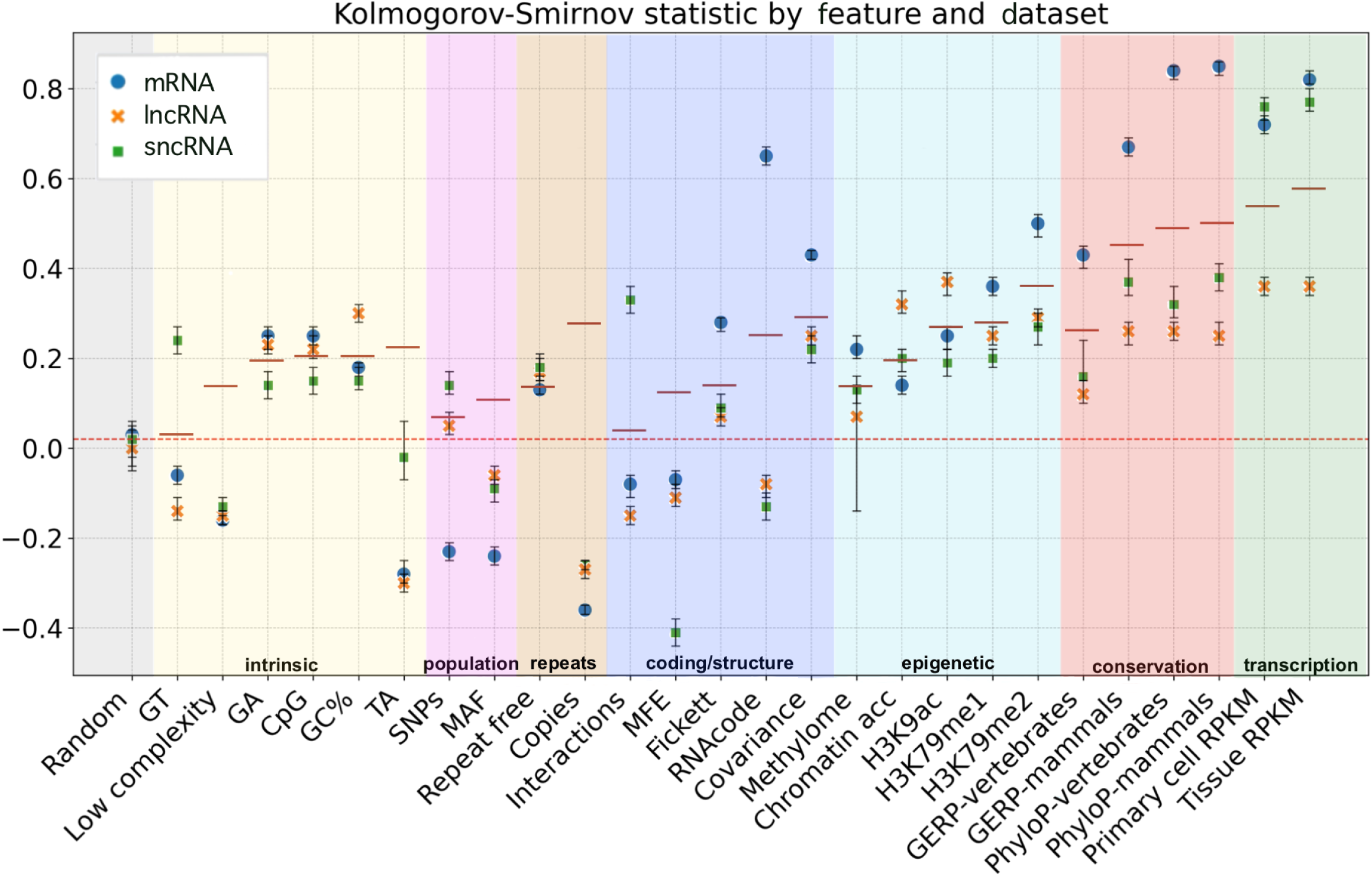
Kolmogorov-Smirnov statistics by feature and dataset. Signed KS statistics comparing functional sequences to the negative control for each feature and dataset. Each point represents the statistic with the 95% confidence intervals, calculated using bootstrapping (1000 replicates). The horizontal dashed red line indicates the value for the Random feature, and background shading the features type. The solid horizontal red lines correspond to the unsigned KS statistic for all three dataset combined (allRNA).

### All features in context

As shown in Figure 1, the most important features that clearly differentiate the different functional genes types and their corresponding negative controls are expression (tissue and primary cell RPKM) and evolutionary conservation (PhyloP, GERP). These two are followed by chromatin state (H3K79me2) and repeat association (copies). These features show the strongest effects across all three main gene types, protein-coding (mRNA), short non-coding RNA (sncRNA) and long non-coding RNA (lncRNA).

Further discriminators of particular note are that coding potential (RNAcode) and secondary structural signatures (Covariance) are key important discriminators for protein coding genes; while secondary structure (MFE) for sncRNA genes and chromatin state (Chromatin) for lncRNA genes distinguish genic from intergenic regions (Supplementary Figure S4).

### Transcription

Transcription is an essential step for the expression of both protein-coding and non-coding genes (Vannini and Cramer 2012; Poliseno et al. 2024). While all genes are transcribed, it is a logical fallacy (affirming the consequent) to assume that all transcribed regions are genes, due to the possibility of transcriptional noise (Raser and O’Shea 2005; Palazzo and Lee 2015; Doolittle and Brunet 2017; Doolittle 2018). To assess transcription signals, we use maximum RPKM (reads per kilobase of transcript per million mapped reads) to summarize transcription across ENCODE Total and Small RNA-Seq experiments from 71 human tissue and 140 human primary-cell samples (ENCODE Project Consortium 2012).

The effect-size for the differences between positive and negative controls for expression features is high (KS statistics range from 0.36-0.82) (Figure 2; Supplementary Table S3), suggesting that genes are highly transcribed compared to negative control sequences. In addition, the robust Z-scores contain rather extreme values [Z ranges from -1.2 to 1.26M for sncRNAs] (Figure 2; Supplementary Table S4; Supplementary Figure S5). This is indicative of extreme differences between the positive and negative sets.

**Figure 2.**
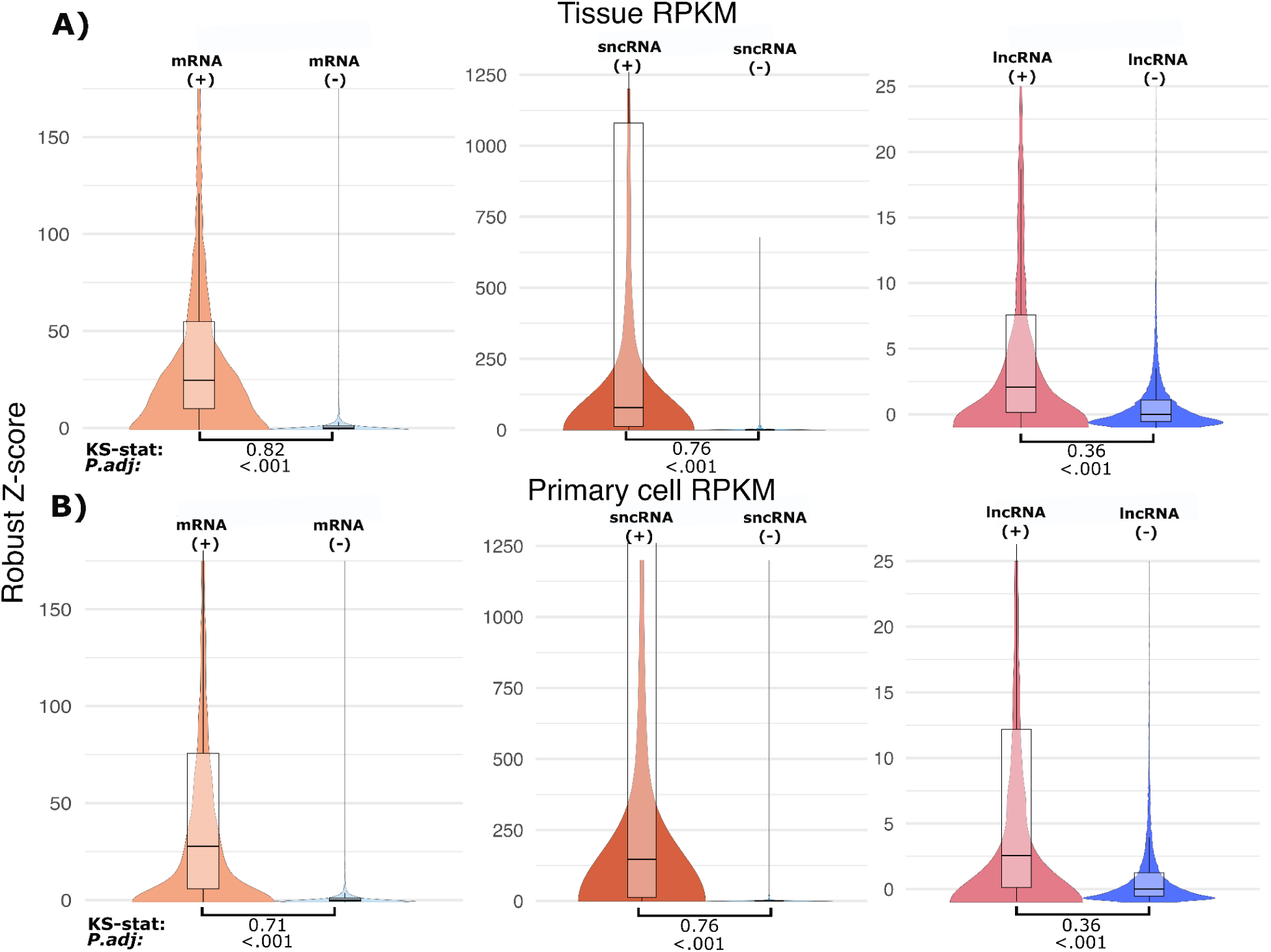
Violin plot and box plots for the distribution of transcriptome-data. A-B) Robust Z-scores for genic and non-genic maximum RPKM across samples (i.e. the highest normalized expression value from RNA-seq data, summarised as reads per kilobase per million (RPKM) mapped reads). Robust Z-scores are computed based upon median, and median-absolute-deviation values from the non-genic controls (**A**) corresponds to tissue samples, (**B**) corresponds to primary cell samples. Signed Kolmogorov-Smirnov statistics and adjusted p-values comparing genic and non-genic controls are displayed under each comparison.

### Conservation

Evolutionary conservation between species is an indicator of function, as negative selection purges variation in these regions due to harmful effects on fitness (Cooper et al. 2005; Nielsen et al. 2007; Christmas et al. 2023). In contrast, non-functional sequences evolve at the neutral rate, which accumulate 1 to 1.5 mutations per 10 megabases per generation (Bergeron et al. 2023). The genome evolution of species with small effective population sizes, such as human, are consequently dominated by drift, and therefore fix many of these mutations (Lynch et al. 2011).

Conservation is assessed here using the phyloP (Pollard et al. 2010) and GERP methods (Cooper et al. 2005; Davydov et al. 2010). We selected datasets that cover both long and short evolutionary distances for each conservation method. For phyloP, we used the short-distance 241-way mammalian dataset (Christmas et al. 2023) and the long-distance 100-way vertebrate dataset from UCSC genome database (Raney et al. 2024). For GERP, we used the short-distance 91-way mammalian dataset and long-distance 63-way amniote vertebrates dataset from Ensembl release 111 (Harrison et al. 2024).

The effect size for differences between positive and negative controls is high for all conservation features (KS-statistics range from 0.12-0.84). This is most pronounced for protein-coding genes: PhyloP 241-way mammals (KS-statistics 0.84) and PhyloP 100-way vertebrates (KS-statistics 0.83) with somewhat lower effects for the GERP scores. These results indicate high evolutionary conservation for both protein-coding and non-coding genes (Figure 3; Supplementary Table S3; Supplementary Figure S6). The lncRNAs, in contrast, showed limited, but not insignificant levels of evolutionary conservation (KS-statistics 0.12-0.24), confirming earlier analyses of lncRNA conservation (Eddy 2014; Kutter et al. 2012; Hezroni et al. 2015).

**Figure 3.**
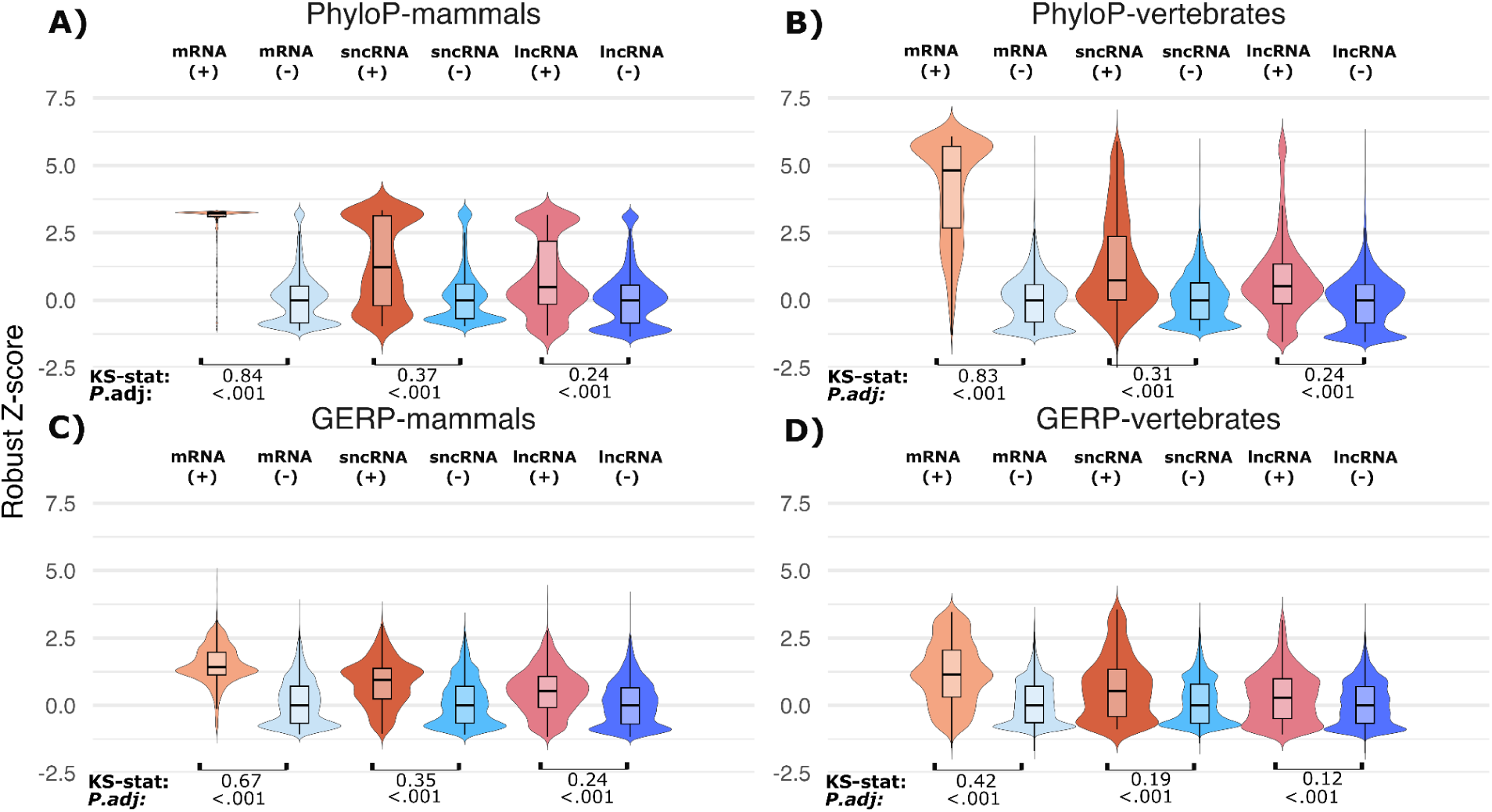
Violin plot and box plots for the distribution of conservation features based on robust z-scores for genic and matched non-genic regions. A&B) PhyloP evolutionary conservation scores for the 241way placental mammal and 100 way vertebrate alignments respectively. **C&D)** GERP evolutionary conservation scores for a 91 way eutherian mammal and 63 way amniote vertebrate alignments respectively. The signed Kolmogorov-Smirnov statistics and adjusted p-values are displayed under each comparison.

The conservation signatures are generally greater for the “short” evolutionary distance mammalian alignments, which may indicate the difficulties of accurately aligning divergent nucleotide sequences from long evolutionary distances (Gardner et al. 2005). Generally the PhyloP scores outperformed GERP for distinguishing genes from non-genic regions.

### Epigenetic features

“Epigenetic” data, such as chromatin accessibility and covalent modifications of DNA or chromatin may be informative of gene activation or suppression (Bannister and Kouzarides 2011; Allis and Jenuwein 2016; Gershman et al. 2022). Regions of accessible chromatin have been associated with active transcription, while compact chromatin indicates transcriptional repression. Open chromatin regions are usually assayed through nuclease or transposon based experiments such as DNase-seq and ATAC-seq (Calo and Wysocka 2013; Chawla et al. 2021). In contrast, cytosine methylation is assayed with bisulphate sequencing and has been associated with transcriptional silencing, which may be indicative of non-genic and regulatory regions (Jones 2012; Greenberg and Bourc’his 2019; Gelfman et al. 2013; Li et al. 2018). Histone modifications regulate DNA accessibility and chromatin structure, providing indications of gene regulatory activity across the genome (ENCODE Project Consortium 2012; ENCODE Project Consortium et al. 2020). In this study histones H3K79me2, H3K79me1 and H3K9ac were selected for optimal discrimination between positive and negative control sequences and minimal redundancy with other modification signatures. We have assessed the epigenetic datasets from ENCODE 3 to determine if this information adds further value for identifying canonical genes (ENCODE Project Consortium et al. 2020).

The effect sizes distinguishing the positive and negative controls with epigenetic features are moderate for chromatin accessibility and methylation (KS statistics: 0.12-0.32) (Figure 4; Supplementary Table S3; Supplementary Figure S7), and comparable to more accessible nucleotide enrichment metrics (e.g., GC%, CpG, TA, GA content; Supplementary Table S3; Supplementary Figure S4). However, both chromatin accessibility and methylation are confounded by sequence composition – GC% and CpG, respectively – with correlations around 0.5, indicating ∼25% of the variance of these genomic datasets is driven by sequence-composition (Supplementary Table S5; Supplementary Figures S1-3). Additionally, both chromatin accessibility and methylation features show weak correlations with transcriptional activity (Tissue RPKM and Primary cell RPKM; mean Spearman’s rho = 0.11 and 0.07) (Supplementary Table S5), consistent with other studies (Kiani et al. 2022).

**Figure 4.**
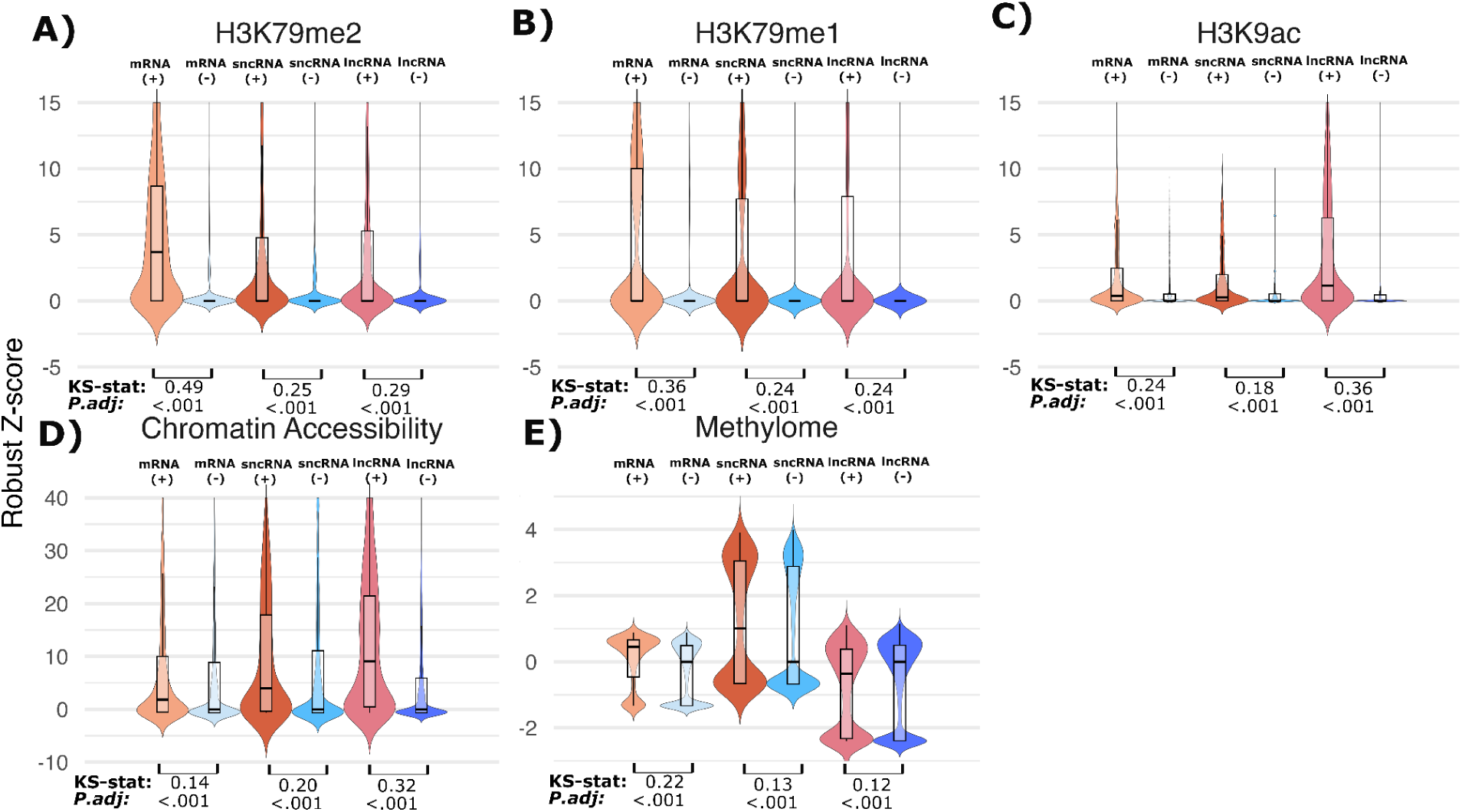
Violin plot and box plots for the distribution of epigenetic features based on robust z-scores for genic and matched non-genic regions. A-C) H3K79me2, H3K79me1 and H3K9ac, maximum signal value for these histone marks from ChIP-seq data. D) Chromatin Accessibility, the maximum signal value for open chromatin regions from DNase-seq and ATAC-seq data. E) Methylome, the average methylation level at CpG sites from bisulphite-sequencing data. All “epigenetic” data was sourced from the ENCODE3 repository. The signed Kolmogorov-Smirnov statistics and adjusted p-values are displayed under each comparison.

In contrast, the effect sizes are comparatively high for the selected histone marks H3K79me1, H3K9ac and H3K79me2 (KS-statistics range from 0.18-0.49) (Figure 4; Supplementary Table S3; Supplementary Figure S8). Each feature has a modest correlation with transcription (Spearman’s rho ranges from 0.06 to 0.35) (Supplementary Table S5), indicating these may be useful orthogonal measures of functional genes.

### mRNA & ncRNA features: structure, interactions & coding potential

#### Protein-coding potential features

A number of methods have been developed for differentiating between coding and non-coding sequences, using metrics such as open-reading frame length, codon usage bias and weighted nucleotide frequencies (Bahiri-Elitzur and Tuller 2021; Champion et al. 2024). While it is unlikely that coding potential can be used to differentiate between ncRNAs and the negative control regions, it is likely that assigned functional protein-coding genes can be classed as “coding” whereas the negative controls will be “non-coding”.

Two independent and accurate metrics were selected to capture coding potential information based on a recent benchmark (Champion et al. 2024): RNAcode uses multiple sequence alignments to calculate a coding potential score based on variation patterns (Washietl et al. 2011), whereas the Fickett score uses compositional bias between codon positions from individual sequences (Fickett and Tung 1992).

As expected, the distributions of RNAcode and Fickett scores for the functional protein coding RNA dataset show large differences from the negative controls (KS-statistics 0.65 and 0.28 respectively), suggesting that it is likely associated with protein-coding functionality, while the ncRNAs effect-sizes were modest (KS-statistics ranged from -0.13 to 0.09) (Figure 5A&B; Supplementary Table S3).

**Figure 5.**
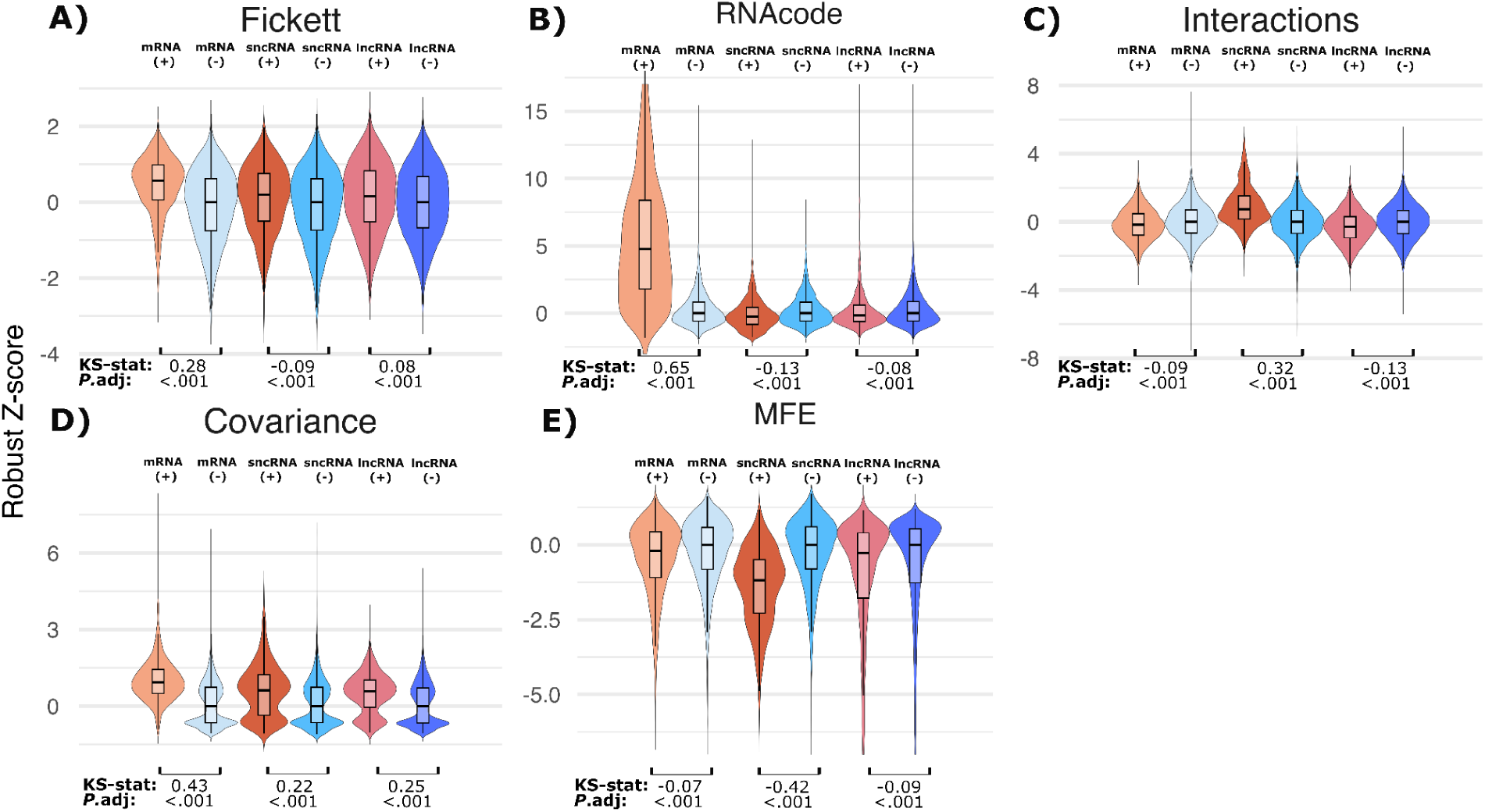
Violin plot and box plots for the distribution of mRNA and ncRNA specific features based on robust z-scores for genic and matched non-genic regions. A&B) Fickett and RNAcode, coding potential based on a measure of nucleotide composition bias and positional asymmetry/ or evolutionary signatures of purifying selection on synonymous sites within multiple sequence alignments, respectively. C) Interactions, average interaction free energies between sequences and 34 abundant human ncRNAs from RNAcentral. D) Covariance, measure of evolutionary pressure to maintain RNA base-pair interactions, reflecting structural constraints. E) MFE, lowest thermodynamic energy of the most stable predicted RNA secondary structure. Signed Kolmogorov-Smirnov statistics and adjusted p-values are displayed under each comparison.

#### RNA structure features

Many functional ncRNAs maintain conserved RNA secondary structures that can be detected with pairwise covariation statistics in sequence alignments (Kertesz et al. 2010; Li et al. 2012; Ding et al. 2014; Rivas et al. 2017; Chiu and Kolodziejczak 1991). A nearest neighbour model of free energy of RNA has been widely used to infer stable secondary structures, however robust controls are required to correct for C+G content, dinucleotide and length to control for confounding factors (Workman and Krogh 1999; Rivas and Eddy 2000).

To capture covarying sites in alignment, which can indicate conserved RNA base-pairs, we used the maximum G-test covariance score implemented in R-scape (Rivas et al. 2017). The effect size for covariance on the difference between functional and negative controls is large for protein and lncRNAs (KS-statistic 0.43 and 0.25 respectively), while the sncRNAs have a more modest difference (KS-statistic 0.22) (Figure 5D; Supplementary Table S3). This finding is unexpected because, outside a limited number of regulatory elements, protein-coding transcripts are not generally thought to preserve long-range RNA secondary structure through compensatory substitutions (Katz and Burge 2003; Rivas et al. 2017; Rivas 2020). Similarly, most lncRNAs show little convincing evidence of conserved base pairing across species (Ulitsky 2016; Haerty and Ponting 2014; Palazzo and Lee 2015). The more modest signal observed for short ncRNAs may reflect the prevalence of classes such as microRNAs and C/D box snoRNAs, where function is frequently mediated by short guide or seed regions and protein interactions, yielding limited detectable covariation in alignments (Brennecke et al. 2005; Gardner et al. 2010; Rivas 2020).

Additionally, because some ncRNAs form stable RNA secondary structures with a low free energy, we estimate the minimum free energy (MFE) of optimal structures for each control sequence with RNAfold (Lorenz et al. 2011). We observed a large effect for the short ncRNAs (KS-statistic -0.42), while the effect sizes were small for the mRNA and lncRNAs as expected (KS-statistics -0.07 and -0.09 respectively) (Figure 5E; Supplementary Table S3).

#### RNA:RNA interactions features

While RNA-RNA interactions play key regulatory roles, there is also evidence of selection against stochastic interactions, leading to widespread interaction avoidance signals in non-coding RNAs (Bhandari et al. 2021; Umu et al. 2016). To quantify this, we used RNAup (Mückstein et al. 2008) to compute RNA-RNA binding free energies between each sequence and a set of 34 abundant, conserved ncRNAs (listed in Supplementary Table S6).

Short ncRNAs have higher Z-scores, suggesting that excess stable RNA-RNA interactions are selected against. We further assessed the strongest sncRNA interactions and found that snoRNA-tRNA interactions are the most common (Supplementary Table S6). This implies that these interactions are not driven by sequence homology, and may further support “orphan” snoRNA directed tRNA modifications (Zhang et al. 2023) (Supplementary Table S6). The interaction excesses or avoidance signals for protein and lncRNA were comparatively modest (KS-statistics -0.09 and -0.13 respectively, Supplementary Table S3) (Figure 5C; Supplementary Table S3).

##### Repeat associations

Repeat elements such as transposable elements, LINEs, SINEs and endogenous retroviruses drive major variations in genome sizes (Osmanski et al. 2023). The bulk of repeats are inactive, and evolve at approximately the neutral rate of evolution (Arkhipova 2018). Transposon mutagenesis techniques such as tnSeq can aid the identification of genomic regions required for survival, and has been well used in bacterial genomics research (Hutchison et al. 1999; Cain et al. 2020). This work inspired a “repeat-free” screen of the human genome, which successfully identified portions of the genome that resist transposon insertions (Simons et al. 2006). We have modeled the “repeat-free” method by determining the combined distance to the nearest upstream and downstream non-overlapping repeat annotation. These include transposable elements, satellite repeats, and pseudogenes from the Dfam database (Hubley et al. 2016; Storer et al. 2021). We excluded repeats overlapping gene annotations as many functional short non-coding RNA genes are classed as repeats by sequence similarity-based annotation tools (e.g.tRNA-derived SINES and the SRP RNA-derived Alu repeats) (Sun et al. 2007; Kriegs et al. 2007).

The effect-size for the differences between positive and negative controls of the repeat-free regions is modest. The signed D statistics from Kolmogorov-Smirnov test are 0.13, 0.17 and 0.16 for proteins, short ncRNAs and lncRNAs respectively (Figure 6A; Supplementary Table S3).

**Figure 6.**
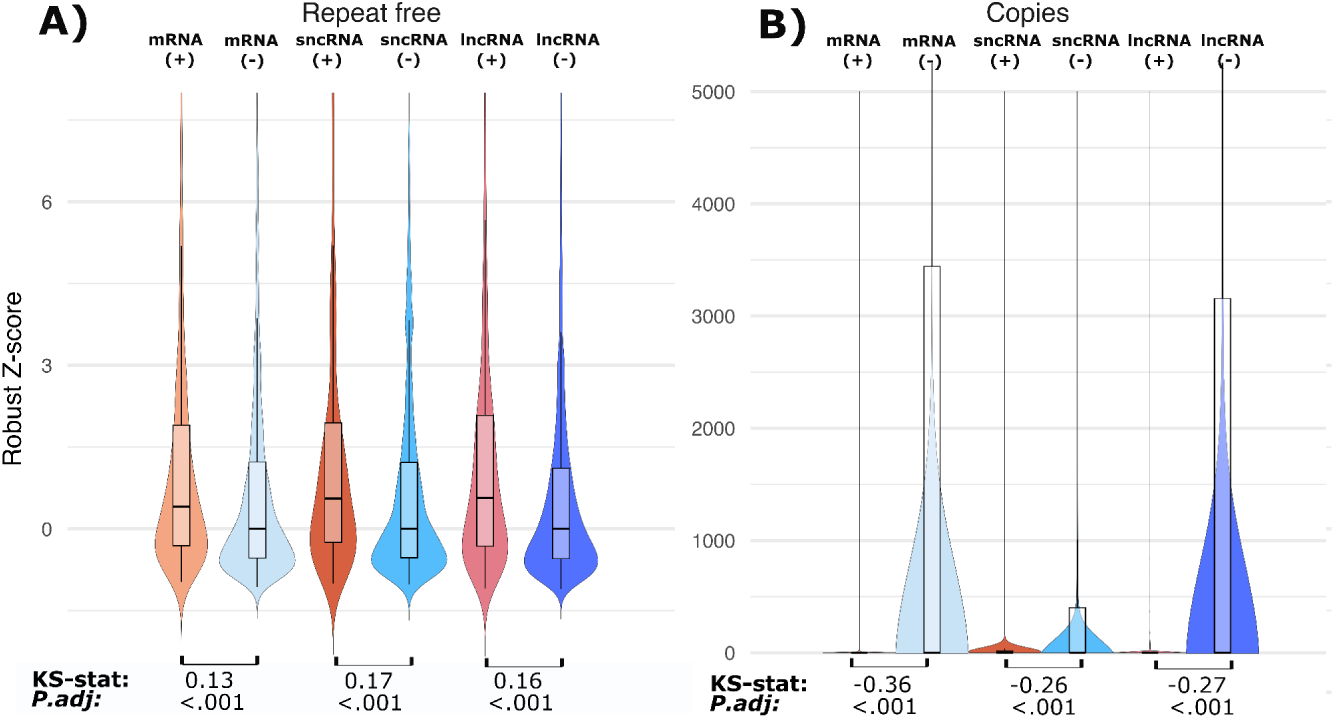
Violin plot and box plots for the distribution of repeat association features based on robust z-scores for genic and matched non-genic regions. A) Repeat free, combined distance to nearest up and downstream repeat elements (SINE, LINE, etc) that does not overlap each control sequence. B) Copies is the genomic copy number of each analyzed region as determined by MMseqs2 searches of the GRCh38 genome. Signed Kolmogorov-Smirnov statistics and adjusted p-values are displayed under each comparison.

We also consider the genomic copy number of each control. Duplicated genes may be more likely to be functional compared to single-copy genes, suggesting that genomic copy-number could be a potential feature of function (Ohno 1970; Bergthorsson et al. 2007; Mihajlovic et al. 2025). Alternatively, regions that have very high copy-number may indicate pseudogenised transposable elements (Osmanski et al. 2023). Either way, copy-number is worth considering as a potential indicator of function.

The copy number results show markedly different distributions between the positive and negative control datasets. Again, the p-values are near zero and the signed D statistics from Kolmogorov-Smirnov test are -0.36, -0.26, -0.27 for proteins, short ncRNAs and lncRNAs respectively. This indicates that genes have far fewer genomic copies than negative control datasets which have high numbers of repeats (Figure 6B; Supplementary Table S3; Supplementary Figure S9-10).

Measures based on sequence conservation (PhyloP, GERP, RNAcode, covariation) show strong correlations with genomic copy number (Spearman’s rho = 0.42-0.62; Supplementary Table S5). While this may partly reflect confounding by sequence composition or complexity, the correlations of these potential confounders with copy-number is weak (Spearman’s rho = 0.01-0.29; Supplementary Table S5).

##### Population variation

Genetic variation between individuals is shaped by selection on function: functionally important regions tend to carry less variation due to purifying selection, while non-functional regions accumulate variants more freely (Vitti et al. 2013). To assess this, we use data from gnomAD v4 (Chen et al. 2024) to calculate SNP density and the average minor allele frequency (MAF) for each region.

The effect-size for the differences between positive and negative controls with population variation features is low for unsigned KS-statistics (0.05-0.24) (Figure 7A&B; Supplementary Table S3). The results suggest that in general protein-coding sequences have fewer SNPs (KS-statistic -0.23 for SNP density) compared to the negative controls. However, for short ncRNAs there are remarkably higher numbers of SNPs (KS-statistic 0.14 for SNP density) (Figure 7A&C; Supplementary Table S3). We explored the enrichment of SNPs in short ncRNAs further and discovered that a few families of ncRNA were affected by elevated SNP counts: spliceosomal RNAs (snRNA) [0.42-2.74 SNPs/nuc], snaR [0.43-2.39 SNPs/nuc], tRNAs [0.06-2.32 SNPs/nuc] and vault RNA [0.34-0.92 SNPs/nuc]. Note that in the absence of INDELs, 3.0 SNPs/nuc is the theoretical maximum (Figure 7C, Supplementary Table S7).

**Figure 7.**
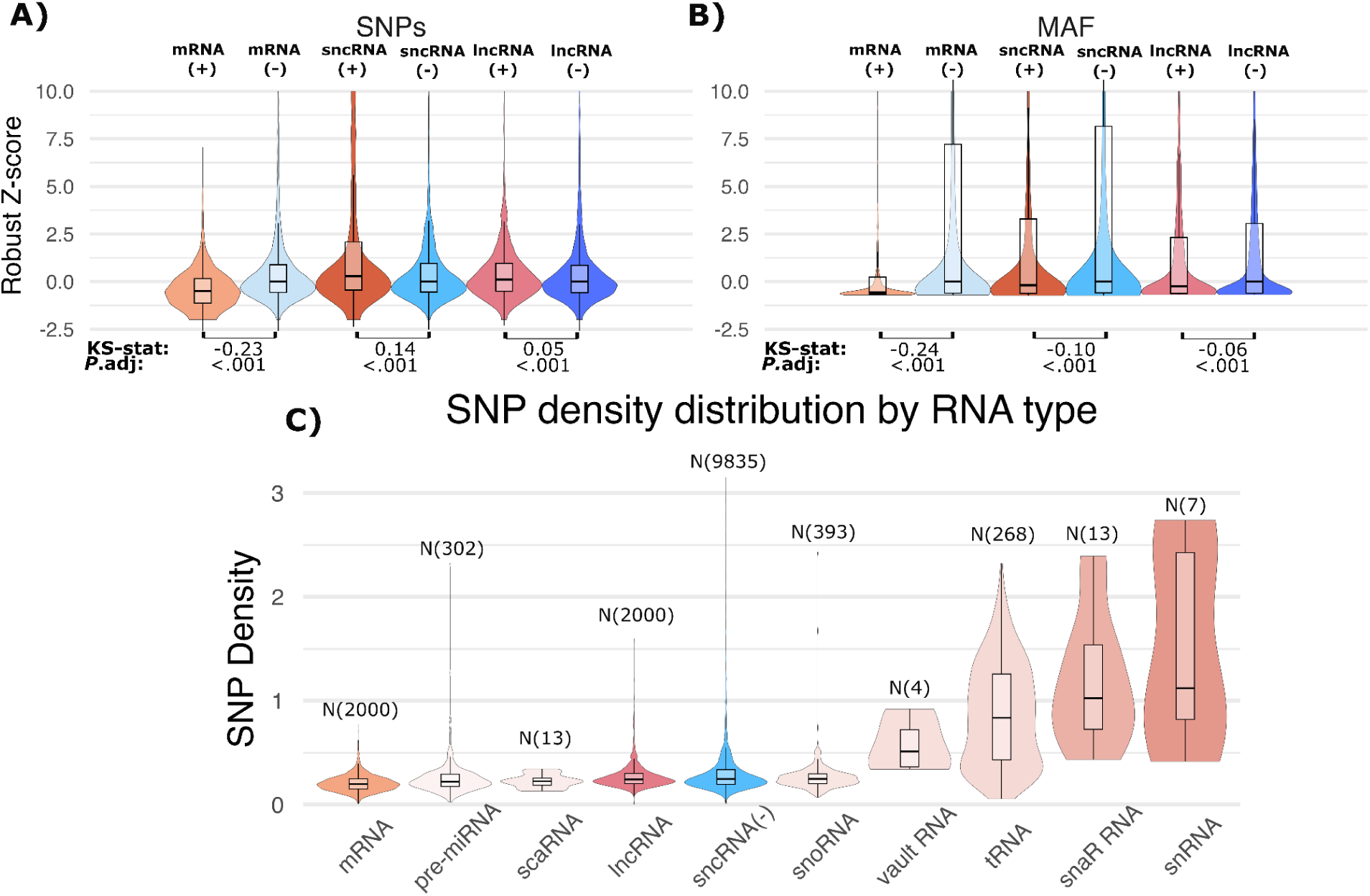
Violin plot and box plots for the distribution of population variation features based on robust z-scores for genic and matched non-genic regions. A) SNP density, measured by the number of SNPs/sequence length. B) MAF is the mean minor allele frequency across all SNPs in a region, where the minor allele frequency is the population frequency of the least common allele at each variant site. C) The SNP density distribution categorized by small non coding RNA types used in this study. Signed Kolmogorov-Smirnov statistics and corresponding adjusted p-values are displayed under each comparison.

Due to the possibility of linkage disequilibrium influencing the SNP statistics, we checked for an association between the distances of each negative control from corresponding positive control (gene). We found the relationships between distance and SNP statistics (e.g. density) to be minimal [Spearman’s rho of -0.01, -0.03 and -0.01 for mRNA, sncRNA and lncRNA genes respectively] (Supplementary Figure S11; Supplementary Table S8).

SNP density is associated with CpG, Low Complexity, GC% and copy-number, with mean absolute Spearman’s rho values of 0.37, 0.33, 0.28 and 0.23, respectively (Supplementary Figures S1-3; Supplementary Table S5). Many of these genomic features may be associated with an increased mutability and mis-annotation due to methylations, polymerase slippage and segmental duplicates (Li 2014; Cooper and Krawczak 1990; Meunier and Duret 2004).

##### Intrinsic sequence features

Intrinsic sequence features are the local components of a sequence. In this work, we include commonly used metrics such as C+G content, the proportion of dinucleotide frequencies (relative to expectations from mononucleotide contents), and the presence of low complexity sequences, which capture simple repeats.

#### Nucleotide statistics

C+G-rich regions have been linked to both protein-coding and non-coding RNA genes (Lander et al. 2001; Haerty and Ponting 2015; Christmas et al. 2025). While among the 16 possible DNA dinucleotides, CpG sites are prone to methylation and subsequent TpG conversion (Medvedeva et al. 2010). Dinucleotides like TA may be underrepresented in genes due to a the suppression of cross-talk with cellular processes such as splice site recognition, AU-rich RNA degradation motifs, “TATA” motif, and premature stop codons (Akinyi and Frilander 2021; Olthof et al. 2024; Barreau et al. 2005; Karlin and Mrázek 1997). Likewise, GT and AG dinucleotides may be avoided due to their roles in splice site identification (Olthof et al. 2024). Further differences in dinucleotide composition may be driven by codon bias or RNA folding and interaction requirements (Parvathy et al. 2022; Su et al. 2024; Umu et al. 2016).

**Low complexity regions** (LCRs), which consist of repetitive or biased subsequences, deviate from random sequence norms (Enright et al. 2023). Within coding sequences, LCRs are thought to evolve neutrally due to low conservation and size variation, driven by polymerase slippage and unequal recombination. As non-functional regions, with potentially detrimental effects, LCRs may be selected against in most genic sequences (Enright et al. 2023; Lenz et al. 2014).

The signed D statistics for the GC% are elevated (KS-statistics range from 0.15-0.30), supporting the previous reports for coding and non-coding genes (Lander et al. 2001; Haerty and Ponting 2015). Meanwhile, the signed D statistics for low complexity density results are low (-0.12 to -0.16), indicating that the coding and non-coding genic sequences have a higher complexity (fewer repetitive sequences) compared to negative control regions, as expected. For the dinucleotide compositions the unsigned D statistics range from 0.03 to 0.30. The most marked differences can be observed for CpG (increased), GA (increased) and TA (decreased) indicating significant shifts of dinucleotides in positive controls compared to negative controls dataset across all three gene types. In contrast, the dinucleotide GT is enriched in the sncRNAs. This is notable as GT is not associated with any known snoRNA or tRNA motif (Figure 8; Supplementary Table S3).

**Figure 8.**
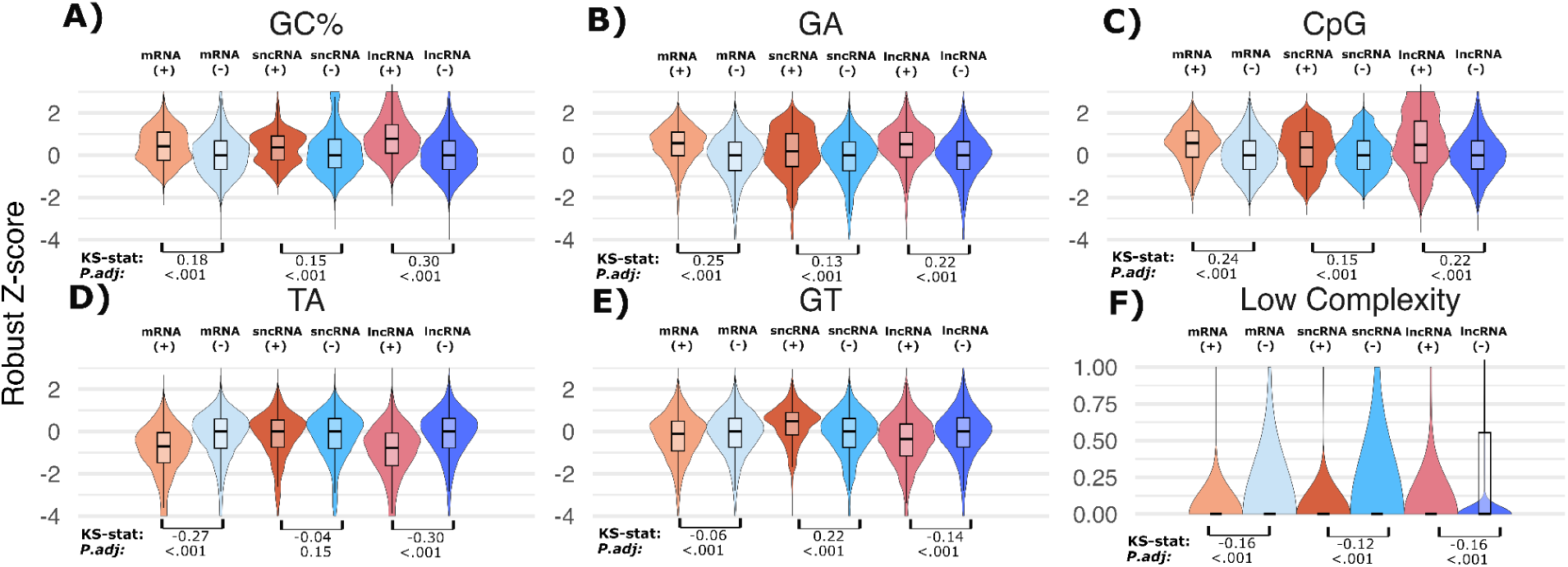
Violin plot and box plots for the distribution of intrinsic sequence features based on robust z-scores for genic and matched non-genic regions. A) GC%, the C+G proportion for each sequence. B-H) selected log-odds of dinucleotides proportions relative to mononucleotide frequencies for GA, CpG, TA, and GT respectively. I) Low Complexity, the proportion of sequence not classed as “low complexity” by the DUST annotation tool. The signed Kolmogorov-Smirnov statistics and adjusted p-values are displayed under each comparison.

## Discussion

In genetics, “function” is interpreted in two main ways. The causal role view considers any biochemical activity – like transcription – as functional. In contrast, the selected effect view requires both biochemical activity and evidence of evolutionary selection, acknowledging that some cellular activity may be mere noise (Eddy 2012; Doolittle and Brunet 2017; Graur 2017). Our analysis is based on the premise that genic exons are distinguishable from intergenic regions with genome-wide signals. By examining a broad range of features, we systematically assessed the relative power of different indicators for separating the genic from non-genic regions. **Transcriptional activity** (RPKM) emerged as the strongest predictor – especially for protein-coding and short ncRNAs, while **evolutionary conservation** was the next most informative, particularly for coding regions. Conservation, however, was generally a weaker signal for ncRNAs (Figures 1–3; Supplementary Fig S4).

Together, these results show that combining **biochemical activity** and **evolutionary selection** offers a robust framework for identifying functional genomic regions. In addition, our feature-based analysis strongly supports the “**selected effect**” model of function, where both signal and selection are needed to reliably distinguish genuine genes from genomic noise (Figure 1; Supplementary Table S3).

Three **histone modifications**, H3K79me1, H3K79me2, and H3K9ac, showed discrimination between gene and control sets comparable to that of transcriptional and conservation signals. However, their strong correlation with transcription and conservation raises questions about whether they offer additional, independent information. In contrast, methylation and chromatin accessibility data showed limited ability to distinguish functional regions. Given this, annotation efforts might be better focused on more informative (and confounded) features such as GC% and CpG enrichment (Supplementary Figure S4).

**Population variation data**, such as SNP density, performed poorly in distinguishing functional from non-functional regions. For protein-coding genes, their performance was comparable to basic sequence features like GC% and CpG content, which are useful but far from definitive. Strikingly, some short ncRNA classes exhibited extremely high SNP densities, in some cases nearing saturation-level (∼3 SNPs per site; Figure 7C). The presence of such excess variation in otherwise conserved genes is unexpected and warrants further investigation to identify the biological or technical causes (Prasetyo and Gardner 2024b).

Among alignment-based features, R-scape’s **covariation score** (max. G-test) was remarkably effective at identifying functional genes, including protein-coding sequences, despite being developed to detect conserved RNA secondary structure (Rivas et al. 2017; Rivas 2020). This unexpected signal may reflect compensatory variation at synonymous sites, a byproduct of the genetic code itself. Similarly, RNAcode’s **coding potential metric** showed strong predictive power for distinguishing coding regions from background based on sequence alignments. In contrast, the **free energy-based features** (e.g. MFE, RNA-RNA interaction scores) were strongest in short ncRNAs, but provided little signal in lncRNAs or protein-coding genes (Supplementary Table S3).

In summary, protein-coding regions are clearly distinguishable from genomic background, supported by strong signals across multiple genomic features (mean unsigned KS = 0.348; Supplementary Table S3). Short non-coding RNAs also show robust functional signatures (mean KS = 0.237; Supplementary Table S3), consistent with their well-characterised roles. In contrast, lncRNAs display weaker and inconsistent signals, with a lower mean KS statistic (mean unsigned KS = 0.201; Supplementary Table S3), suggesting that many lncRNAs are indistinguishable from intergenic (presumed non-functional) regions based on current features. This highlights the need for more stringent criteria or improved annotations for lncRNA function which have historically required remarkably little evidence (e.g. >1 RNA-seq read) (Xu et al. 2017).

### Limitations of the study

#### Annotation Uncertainty

Some positive control annotations may reflect misannotation or overinterpretation of limited evidence. For example, nearly a third of protein-coding genes in the original human genome draft have since been merged or removed (Hatje et al. 2019; Amaral et al. 2023). Similar issues likely affect non-coding annotations, where pseudogenes or spurious transcription may have been over-enthusiastically mislabelled as functional (Prasetyo and Gardner 2024a). Unfortunately, there are fewer opportunities for systematic re-curation of non-coding annotations compared to coding ones. Conversely, some negative controls may include unannotated functional elements, although both types of errors are expected to be rare given the decades of intensive curation and discovery efforts in annotating the human genome.

#### Transcriptome Coverage and Circularity

While we observe strong associations between gene expression, chromatin marks, and annotated genes, our RNA-seq datasets - though extensive (211 samples) - may miss tissue-specific or condition-specific transcripts. As a result, “unexpressed” regions may not truly be inactive. The discovery of many lncRNAs using RNA-seq and chromatin data (Xu et al. 2017), also introduces potential circularity, inflating associations between those features and lncRNA annotations.

#### Negative Control Design

Our choice of negative control sequences, intergenic regions matched by location and sequence properties, may not be ideal. Alternative approaches, such as synthetic random sequences or shuffled chromosomes, are rarely available and not suitable for comparing evolutionary or population variation signals (Eddy 2013; Camellato et al. 2024).

#### Confounding Variables

Some confounding factors may have been overlooked. We addressed the most obvious by matching for length and local GC/CpG content, but deeper latent biases may still affect certain features.

#### Class Imbalances

To reflect real genome composition, we deliberately included more negative than positive controls, consistent with prior estimates of functional genome fractions (Galeota-Sprung et al. 2020; Christmas et al. 2023). However, within positive sets, imbalance between RNA classes posed a challenge: sncRNAs are dominated by miRNAs (30.2%), snoRNAs (39.3%), and tRNAs (26.8%), requiring downsampling to avoid over-representation in downstream analyses.

## Materials and Methods

We will next describe the positive and negative control annotations we selected, the genomic features used for analysis, and the analytical methods employed to identify features associated with functionality.

### Retrieval of functional genes

We designated sequences as “functional” based on annotations from the HUGO Gene Nomenclature Committee (HGNC) database, which links each gene to documented evidence of function (Braschi et al. 2019). Because named genes undergo multiple rounds of expert curation, we considered them more likely to be functional.

We retrieved protein-coding sequence IDs from HGNC and filtered out withdrawn entries or those with multiple RefSeq IDs (Braschi et al. 2019). Using these RefSeq IDs, we extracted chromosome coordinates and corresponding sequences from the GRCh38.p14 genome (O’Leary et al. 2016).

To include functional non-coding RNAs, we sourced sncRNAs and multi-exonic lncRNAs from RNAcentral (release 22.0), including only those with assigned HGNC ID. We downsampled precursor miRNAs, which are overrepresented in RNAcentral, to balance the representation of different sncRNAs types (Freyhult et al. 2005; The RNAcentral Consortium 2019). Our sncRNA dataset includes small nucleolar RNAs (snoRNAs) (39%), transfer RNAs (tRNAs) (27%), small nuclear RNAs (snRNAs) (0.7%), microRNAs (miRNAs) (30%) and other less represented groups such as vault, sca and snaR RNAs (3%), with precursor miRNAs added to complete the 1,000-sequence set.

We excluded chromosome Y and mitochondrial DNA due to the former’s excess low complexity sequence, and the latter’s different selection pressures (Quintana-Murci and Fellous 2001). We also excluded ribosomal RNAs because of their large sequence length, unusual expression and sequence conservation patterns, and uncertain placement in the reference human genome (Agrawal and Ganley 2018).

From each functional gene category, we randomly selected up to 1,000 sequences. For protein-coding genes, we created two datasets: one for exon two and another for exon three, focusing on internal exons and avoiding untranslated regions and promoters, which are subject to different evolutionary constraints. For lncRNA genes, we generated separated datasets for exon one and exon two, due to the limited number of multi-exonic annotations.

To minimize length-dependent effects and ensure computational feasibility- given that some of the algorithms scale poorly with sequence length-, we filtered sequences by length. We defined the upper and lower limits as the 10th and 90th percentiles of the length distribution of each RNA class: protein-codingRNA (61-271 nucleotides), lncRNA (74-1500 nucleotides), and sncrna (71-142 nucleotides). This approach preserved statistical robustness while maintaining manageable input sizes for downstream analyses.

### Negative control dataset

To generate the negative control datasets, we sampled up to 10 negative control sequences per functional sequence from the GRCh38.p14 human genome (O’Leary et al. 2016). We filtered these sequences to exclude any overlaps with annotated genes. Using Bedtools 2.29.0, we removed controls that intersect with RNAcentral and GENCODE annotations (Quinlan 2014; Haeussler et al. 2019; Harrow et al. 2012). We selected these databases for their extensive annotation coverage, significant overlap with other resources (GENCODE v45), and the high number of unique entries provided by RNAcentral (Supplementary Figure S12).

We chose control coordinates from regions located approximately 1, 10, 100, 1000, and 5000 kb upstream and downstream of each functional sequence. These distances allowed us to sample broadly across “non-genic” regions and assess the influence of local genomic signatures (Elhaik and Graur 2014; Chor et al. 2009; Costantini et al. 2009) (Supplementary Figure S13).

To control for sequence length bias, we matched each negative control sequence to the length to its corresponding functional sequence. We excluded sequences with more than 5% ambiguity (i.e. “N”).

We deliberately set the functional-to-control ratio at approximately 1:10, reflecting recent estimates of the proportion of the genome under functional constraint and negative selection (Christmas et al. 2023). This ratio aligns with our broader goal of applying these features to large genomic regions - or even genome-wide - where most sequences are expected to be non-functional.

### Feature inclusion criteria

Genomic features were included if they met the following criteria: 1. prior evidence of association with gene function; 2. applicable to genome-wide analyses including the negative controls; 3. Informative without redundancy; 4. minimal bias towards known genes (e.g. not homology-based); 5. available for GRCh38; and 6. accessible, installable software where required.

Starting from an initial pool of 98 features, we applied these criteria and excluded 72 that failed to meet one or more requirements. We retained 26 features for downstream analysis (Supplementary Tables S1 & S2) and included a baseline pseudorandom number (“Random”).

### Transcriptome expression features

Transcriptional activity was estimated using ENCODE RNA-seq datasets comprising 71 human tissues and 140 primary cell samples (ENCODE Project Consortium 2012). Libraries were rRNA-depleted; total RNA-seq captured transcripts (≥200 bp), and small RNA-seq targeted transcripts (<200 bp). Dataset accession IDs are provided in the project repository (Schiavinato). Expression levels were quantified as RPKM (Reads Per Kilobase of transcript per Million mapped reads) using SAMtools v1.10 (Li et al. 2009), and the maximum RPKM across all datasets was recorded for each sequence. The maximum RPKM was chosen to capture tissue- or condition-specific expression, which may be obscured by averaging across diverse samples, although this approach may be sensitive to outliers and technical variability.

### Sequence conservation features

We extracted PhyloP scores to analyse sequence conservation from the UCSC hg38 phyloP 100-way and phyloP 241-way bigWig files, which represents multiple alignments of 100 vertebrate species and 241 placental mammals, respectively. To obtain these scores, we used the bigWigSummary command-tool from the UCSC Genome Browser’s ‘kent’ bioinformatic utilities (v385) (Haeussler et al. 2019). We also retrieved GERP (Genomic Evolutionary Rate Profiling) scores for each region using Bedtools 2.29.0, based on the release of 111 Ensembl multiple genome alignments that included 91 eutherian mammals and 63 amniote vertebrates (Cooper et al. 2005; Quinlan 2014; Yates et al. 2020). For each region and method, we used the maximum score as the summary statistic, as this consistently produced the strongest effects in our preliminary evaluation.

### Epigenetic signature features

To assess the contribution of epigenetic features to gene functionality, we obtained processed ENCODE3 datasets (narrowPeak and bedMethyl formats) spanning ∼110 tissues, 91 cell lines, 33 in vitro differentiated cells, and 76 primary cell types (ENCODE Project Consortium et al. 2020) ([CSL STYLE ERROR: reference with no printed form.]). From an initial set of 32 histone modifications, we selected **H3K79me1, H3K79me2, and H3K9ac** using a two-step feature selection procedure: (i) Kolmogorov–Smirnov tests to identify features that best distinguish genic and control regions, followed by (ii) Spearman correlation analysis to retain informative, minimally redundant marks (Supplementary Figure S8). **Chromatin accessibility** and **CpG methylation** were included as additional epigenetic features.

We use an approach that is consistent with the transcriptome analysis above. For each feature, we calculated the maximum signal across all samples and scaled by the degree of overlap between control sequence annotations and the corresponding histone marks or chromatin accessibility regions. I.e.

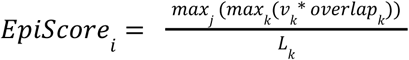

For each control sequence (*EpiScore_i_*), signal values (*v_k_*) are obtained from narrowPeak ENCODE files, which give enrichment values using local background normalization (Feng et al. 2012). The *overlap_k_* value is the overlap length between the control sequence *i* and histone mark or chromatin accessibility region *k*. *L_k_* is the length of control sequence *i*. The max signal is chosen across all *j* tissues and cell lines.

For **methylome** measures, we used a different approach to account for saturated sites, which becomes inevitable given enough samples. Instead, we calculated an average proportion of methylations per CpG site as follows:

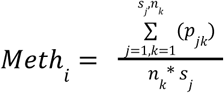

Where *Meth _i_* is the average proportion of CpG methylations per sample per CpG site for sequence *i*. The proportions *p_jk_* are the proportion of methylated sites identified at site *k* in sample *j* (*s_j_*). The number *n_k_* is the number of CpG sites within each control sequence.

### Protein and RNA specific features

We analyzed protein and RNA-specific features using multiple sequence alignments (MSAs), coding potential metrics, RNA structural properties, and RNA:RNA interaction profiles. To obtain MSAs, we extracted alignments from the Zoonomia 241 Placental Mammals dataset (Zoonomia Consortium 2020) using the bigBedToBed command-line tool from UCSC Genome Browser’s ‘kent’ bioinformatic utilities (v385) (Haeussler et al. 2019; Lorenz et al. 2011; Zoonomia Consortium 2020)). We filtered out partial alignments that covered less than 50% of the *Homo sapiens* reference sequence.

We assessed coding potential using CPC2 beta and RNAcode (ViennaRNA 2.4.14), applying either Fickett scores derived from individual sequences, or RNAcode scores calculated from the corresponding MSAs (Kang et al. 2017; Lorenz et al. 2011; Washietl et al. 2011).

To evaluate RNA structure, we used R-scape 1.4.0 to calculate maximum covariance score (G-test) for each region based on the MSA (Rivas et al. 2017). We also applied RNAfold from ViennaRNA 2.4.14 to compute the minimum free energy (MFE) value for each sequence (Lorenz et al. 2011; Bhandari et al. 2019).

To explore RNA-RNA interactions and potential avoidance mechanisms, we used RNAup 2.3.3 to calculate average interaction free energies between each sequence and a curated database of 34 abundant human ncRNA sequences (Supplementary Table S6) from RNAcentral v15, which are excluded from the functional dataset (The RNAcentral Consortium 2019). These ncRNAs are known to interact with a variety of RNAs (Driedonks and Nolte-’t Hoen 2018; Marvin et al. 2011; Yan et al. 2019; Mann et al. 2017).

### Genomic repeat associated features

We estimated the genomic copy number of each sequence by aligning it against the GRCh38.p14 human genome using MMseqs2 v15.6f452 (Hauser et al. 2016). We included all sequence matches with an E-value smaller than 0.001. To calculate the “repeat-free” feature, we used non-redundant hits of repetitive DNA elements in the human genome obtained from Dfam v3.8. These annotations include transposable elements (LINEs, SINEs, LTRs, DNA transposons, other Interspersed repeats), satellite repeats (including TAR), pseudogenes and conserved repeats, identified via profile HMMs.

We identify the closest repetitive element up- and downstream of the sequence using Bedtools 2.29.0 (Quinlan 2014; Hubley et al. 2016). We excluded repeats that overlapped with the control regions as many small ncRNAs are classified as repeats (e.g. tRNAs, spliceosomal RNAs, …). For each sequence, we recorded the sum of the distances to the nearest up and downstream repeats, following a strategy similar to the historic “repeat-free” approach (Simons et al. 2006).

### Population variation features

To assess population variation, we calculated SNP density and average minor allele frequency (MAF) for each genomic region. Variant data was obtained from gnomAD v4 VCF files using Tabix v1.10 (The 1000 Genomes Project Consortium 2015; Li 2011; Karczewski et al. 2020). SNP density is defined as the number of SNPs per base pair, and average MAF is the mean MAF across all SNPs within each sequence. RNA gene types were assigned using RNAcentral, and SNP density distributions were compared across RNA types (Figure 7C).

### Intrinsic sequence features

We calculated G+C content by determining the percentage of guanines and cytosines in each sequence. To assess low complexity regions, we used DUST 1.0.0 (Morgulis et al. 2006) to compute low complexity density across the dataset.

For each dinucleotide, we calculated a log-odds (bitscore) that quantify sequence composition:

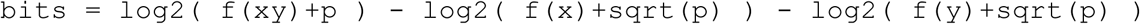

where `f(xy)` is the frequency of dinucleotide `xy`, and `f(x)` and `f(y)` are the frequencies of nucleotides `x` and `y`, respectively. A small pseudocount (`p=0.01`) was added to avoid zero values, particularly in short sequences, and to ensure stable log-ratio estimates.

### Random number

To establish a null baseline, each sequence was assigned a random integer (0-32,767) generated using the Bash $RANDOM function, which produces approximately uniformly distributed pseudo-random values. This feature was included as a control to identify signals no stronger than random expectation (Figure 1).

### Robust z-scores for features

In order to place all genomic features on comparable scales we computed robust z-scores (Rousseeuw and Croux 1993) These also allow comparable assessments of differences between the positive and negative control sequences for each feature. Given the very extreme distributions in our genomic data, we have favored robust Z-scores which use the median and the median absolute deviation (MAD) compared to more sensitive means and standard distributions. The following formula was used (Rousseeuw and Croux 1993):

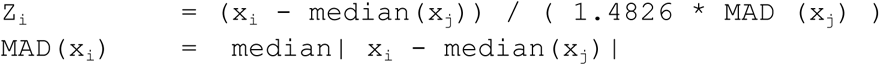

Where x_j_ is the data corresponding to negative control sets for protein-coding, short or long ncRNA. I.e. Each dataset for each gene type is normalized such that the corresponding negative control has a median of zero and a standardized variation. This normalization improves both for visualization – by placing features on comparable scales – and for statistical evaluation across data types. However, despite this normalisation, some panels use independent y-axis scales to accommodate features with highly unequal variance and to allow their distribution to be visualized without excessive compression.

### KS statistics for features

We assessed the direction and absolute difference in distributions between positive and negative control sequences using a signed Kolmogorov-Smirnov (KS) test. We selected this test for its robustness and non-parametric nature, which allowed us to compare distributions without assuming an underlying data distribution (Kini et al. 2025). The KS test’s ability to detect location and shape differences in empirical distributions makes it useful for detecting subtle but significant variations. We used the ks.test and p.adjust functions within the standard R pdackage “stats” v 4.4.1 to assess significance and then p-values were adjusted using BH/FDR correction across all features (number tests were equal to number of features -> 27 in total). To obtain the direction of the stochastic difference between distributions, we computed the cumulative distribution function’s (CDF) maximal and minimal difference and returned the signed value. The confidence intervals were calculated using a bootstrap approach (N=1,000).

## Data Access

All custom scripts, information on downloaded primary data and the generated datasets are available on GitHub: https://github.com/danischiavi/gene-functionality

## Supplementary Information

Supplementary Figures S1-S14

https://github.com/danischiavi/gene-functionality/blob/2024-gene-functionality/docs/Supplementary_figures.pdf

Supplementary Tables S1-S8

https://github.com/danischiavi/gene-functionality/blob/2024-gene-functionality/docs/Supplementary_tables.xlsx

