## Supplementary text and figures S1-S14 for "Genomic indicators of gene function: A systematic assessment of the human genome"

**Supplementary material**

A Spearman correlation matrix was generated to assess correlation among selected features of gene functionality. These plots were created using the R functions ‘ggcorrplot’ from ggcorrplot v.0.1.4.1 package and ‘p.adjust’ from Stats v.4.4.1 package

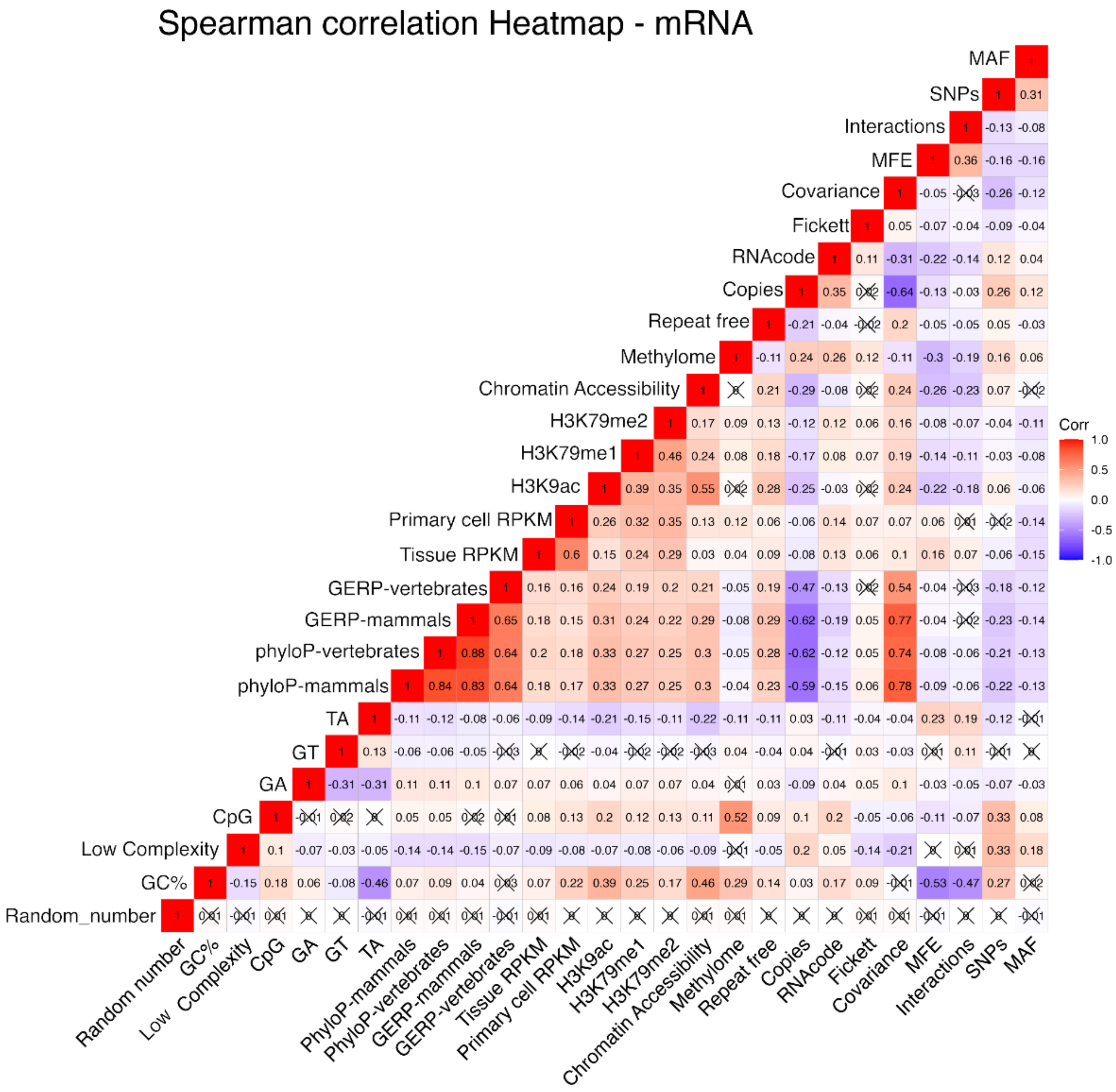


**Supplementary Figure S1. mRNA correlation matrix.** Spearman correlation of functionality predictors for protein-coding sequences. Positively correlated features are displayed in red and negatively correlated features in blue. Spearman’s Rho was used to determine the significance of each correlation, which were corrected for multiple testing using a Bonferroni correction. Non-significant correlations with a p-value > 0.05 have been marked with a cross.

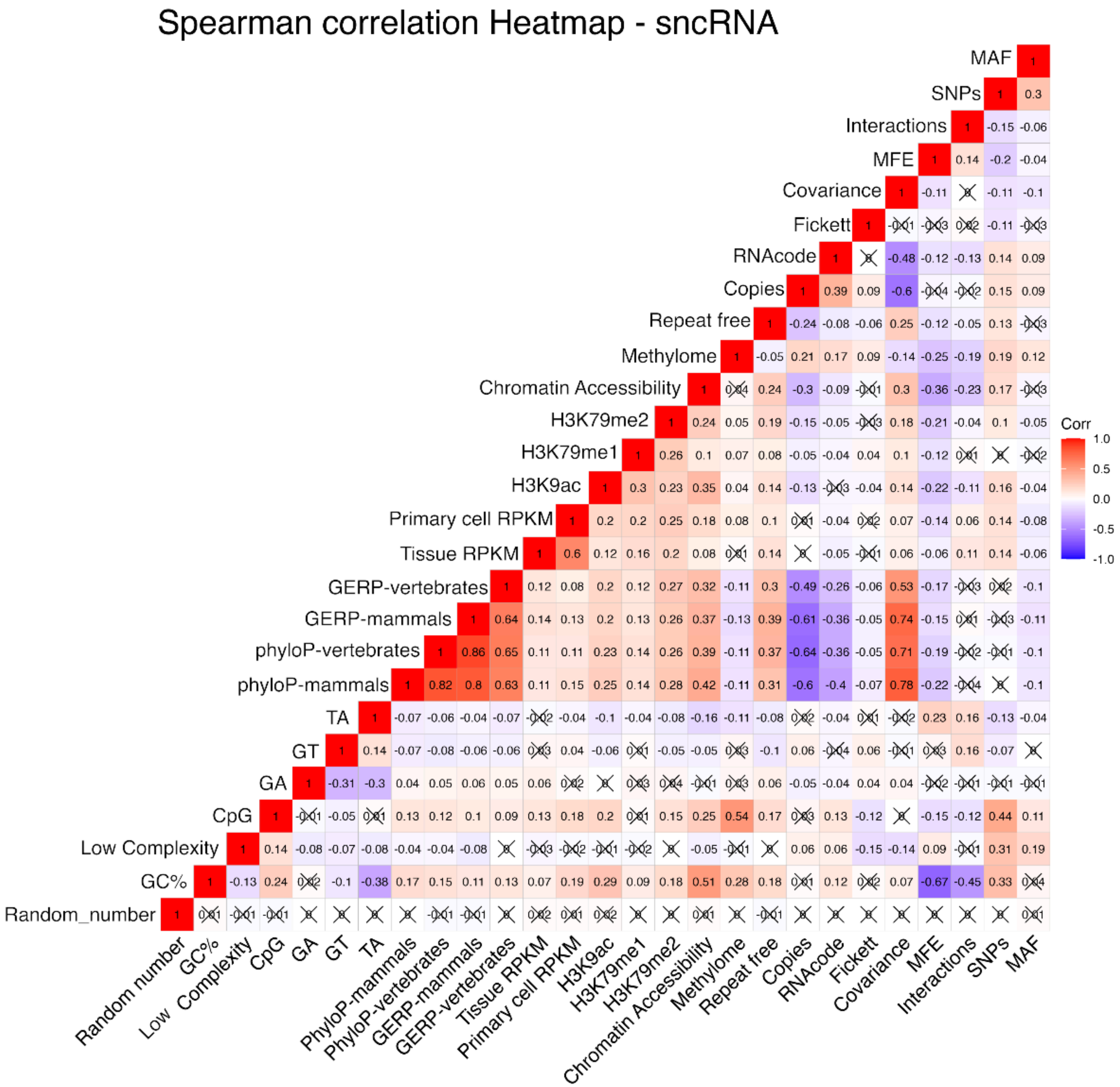


**Supplementary Figure S2. sncRNA correlation matrix.** Spearman correlation of functionality predictors for sncRNA sequences. Positively correlated features are displayed in red and negatively correlated features in blue. Spearman's Rho was used to determine the significance of each correlation, which were corrected for multiple testing using a Bonferroni correction. Non-significant correlations with a p-value > 0.05 have been marked with a cross.

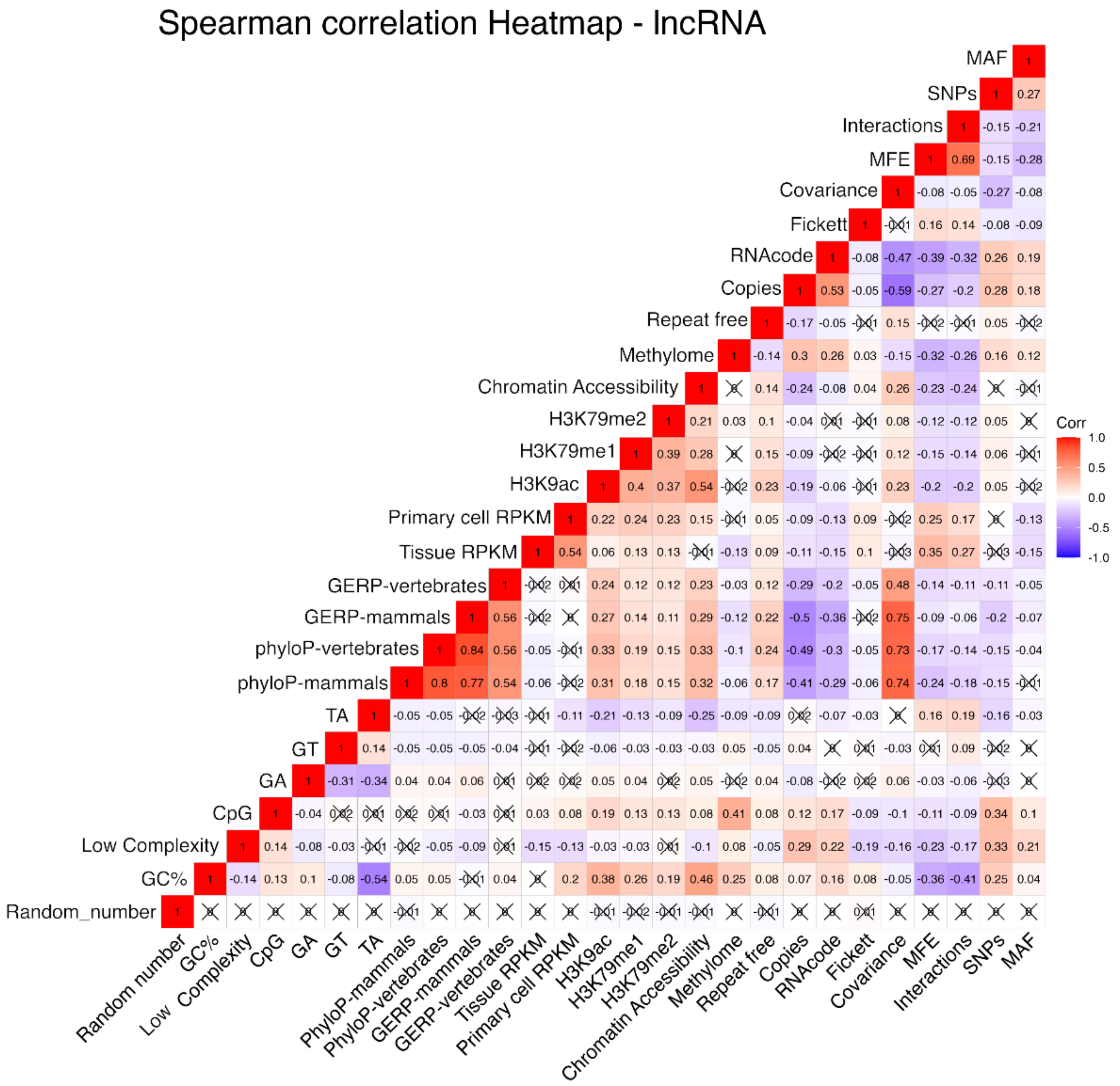


**Supplementary Figure S3. lncRNA correlation matrix.** Spearman correlation of functionality predictors for lncRNA sequences. Positively correlated features are displayed in red and negatively correlated features in blue. Spearman's Rho was used to determine the significance of each correlation, which were corrected for multiple testing using a Bonferroni correction. Non-significant correlations with a p-value > 0.05 have been marked with a cross.

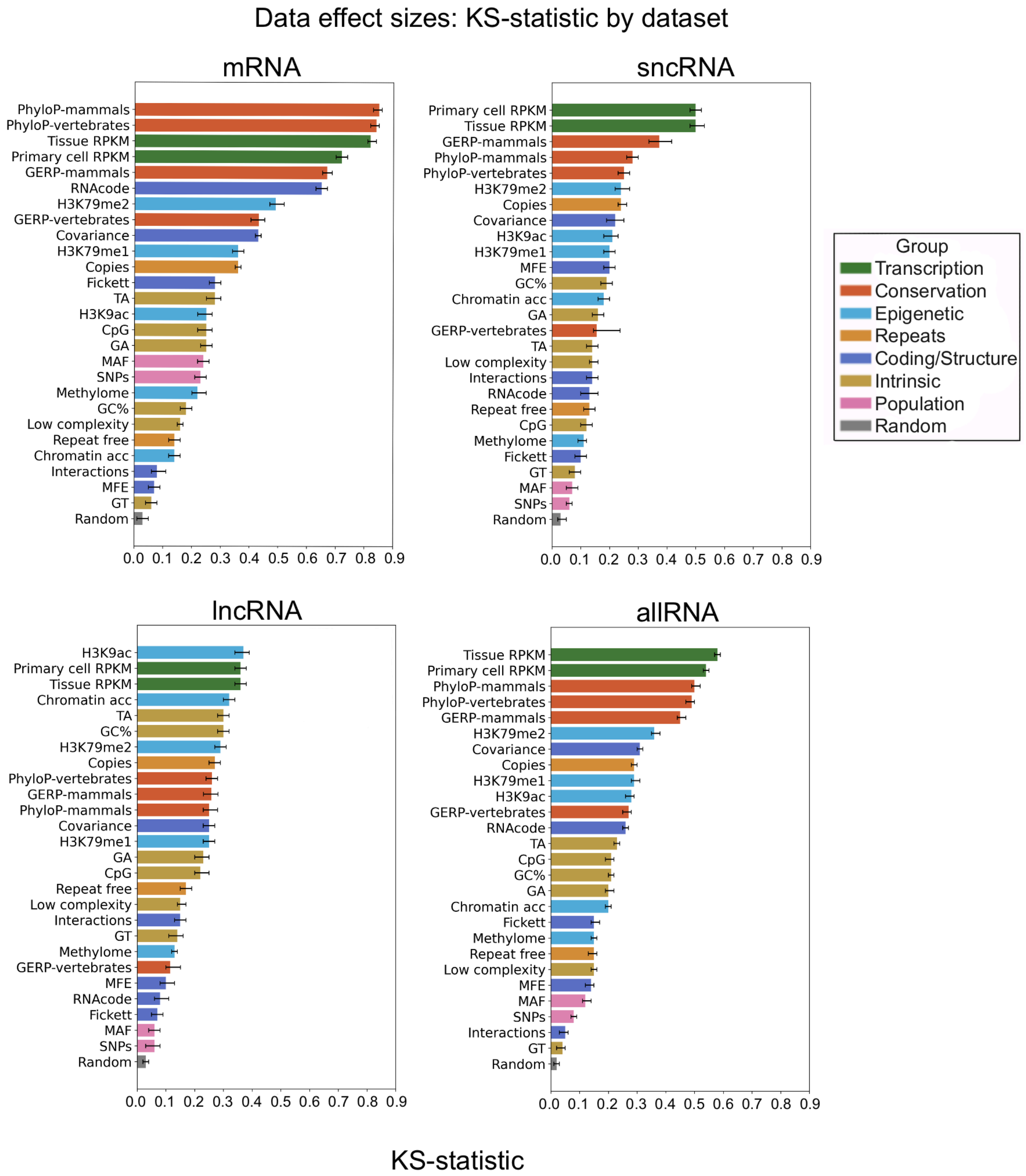


**Supplementary Figure S4. Data effect sizes.** Features ranked by Kolmogorov-Smirnov (KS) statistic. Each point includes 95% confidence intervals, calculated using bootstrapping (1000 replicates). Different colors represent feature classes.

**Genome feature example distributions summaries**

For some of the genome features we illustrate example distributions that are used to generate the robust z-scores violin plots shown in the manuscript. Several features exhibit highly skewed distributions (e.g., primary cell RPKM), which explains the substantial variation in z-scores observed between the positive and negative gene sets.These plots were created using the R functions ‘geom_histogram’, ‘geom_density’, ‘geom_boxplot’ and ‘stat_qq’ from the ggplot2 v3.5.1 package. The distributions for each feature and class can be found in the GitHub repository ([results/Paper figures/Distribution_plots]()).

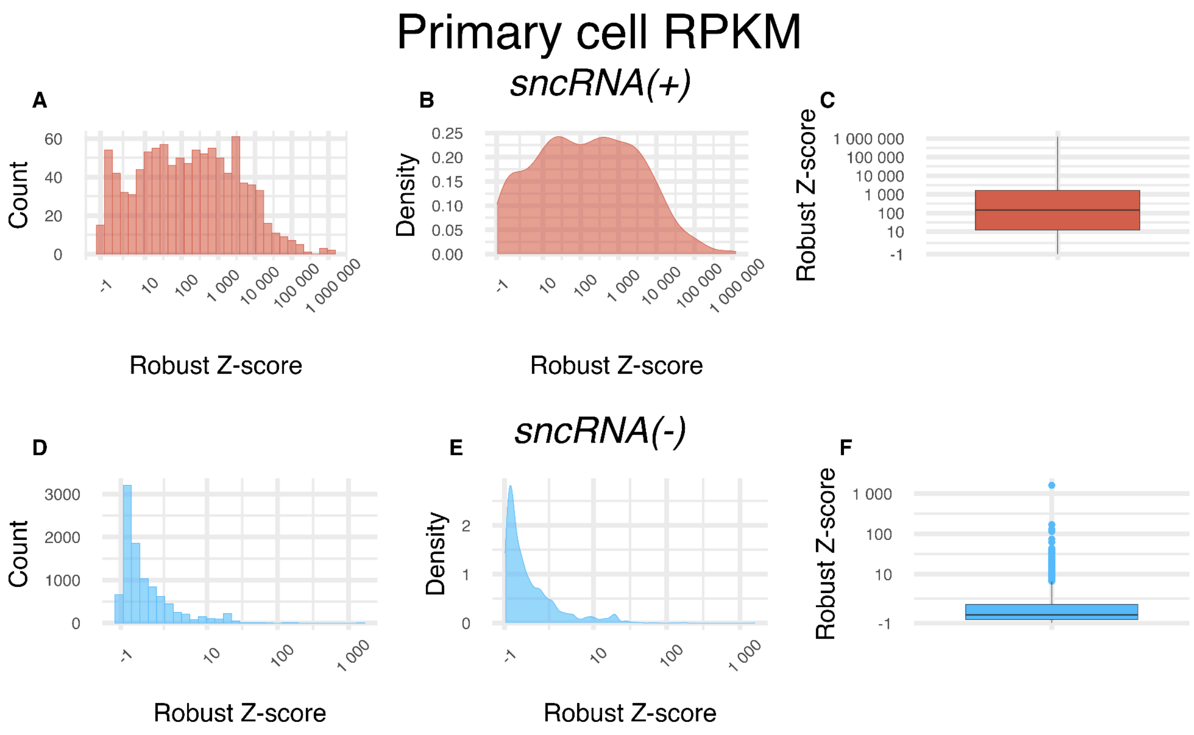


**Supplementary Figure S5. sncRNA primary cell RPKM robust z-scores distributions in log scale.** A-C) Histogram, Density and Boxplot for sncRNA(+), D-F) Histogram, Density and Boxplot for sncRNA(-) all show the robust z-scores for maximum primary cell RPKM values.

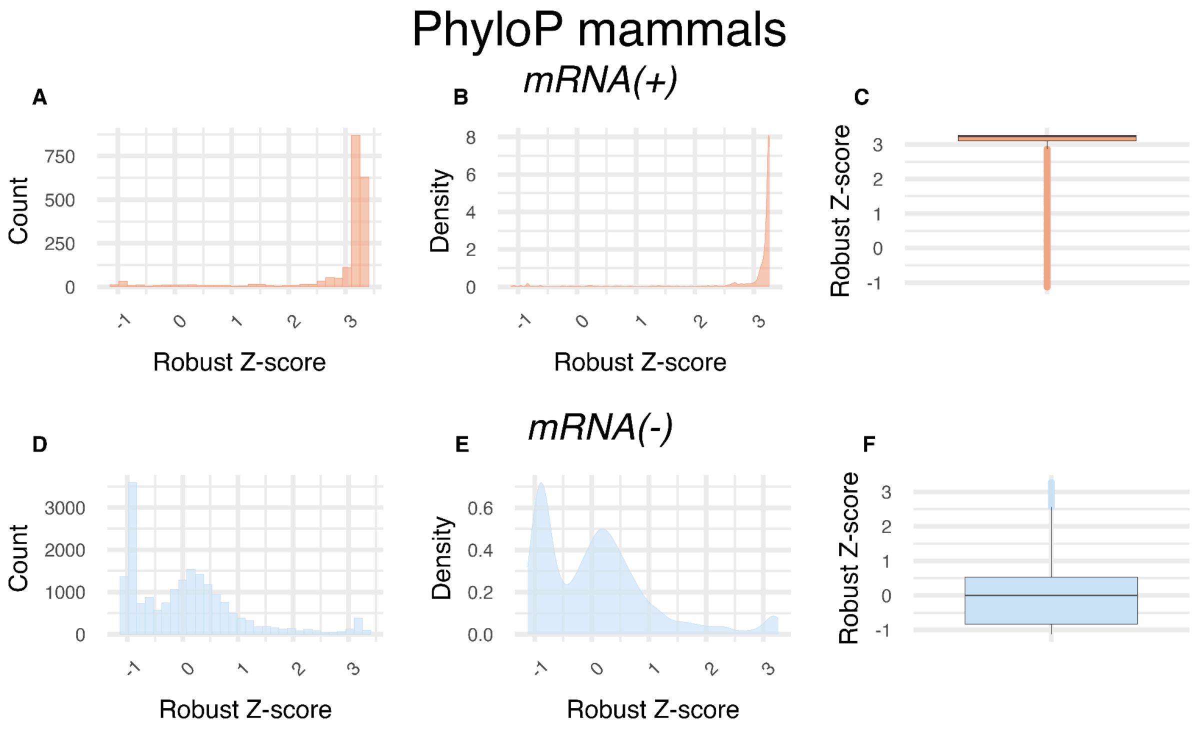


**Supplementary Figure S6. mRNA PhyloP mammals robust z-scores distributions.** A-C) Histogram, Density and Boxplot for mRNA(+), D-F) Histogram, Density and Boxplot for mRNA(-) for robust z-scores generated from PhyloP mammal conservation values.

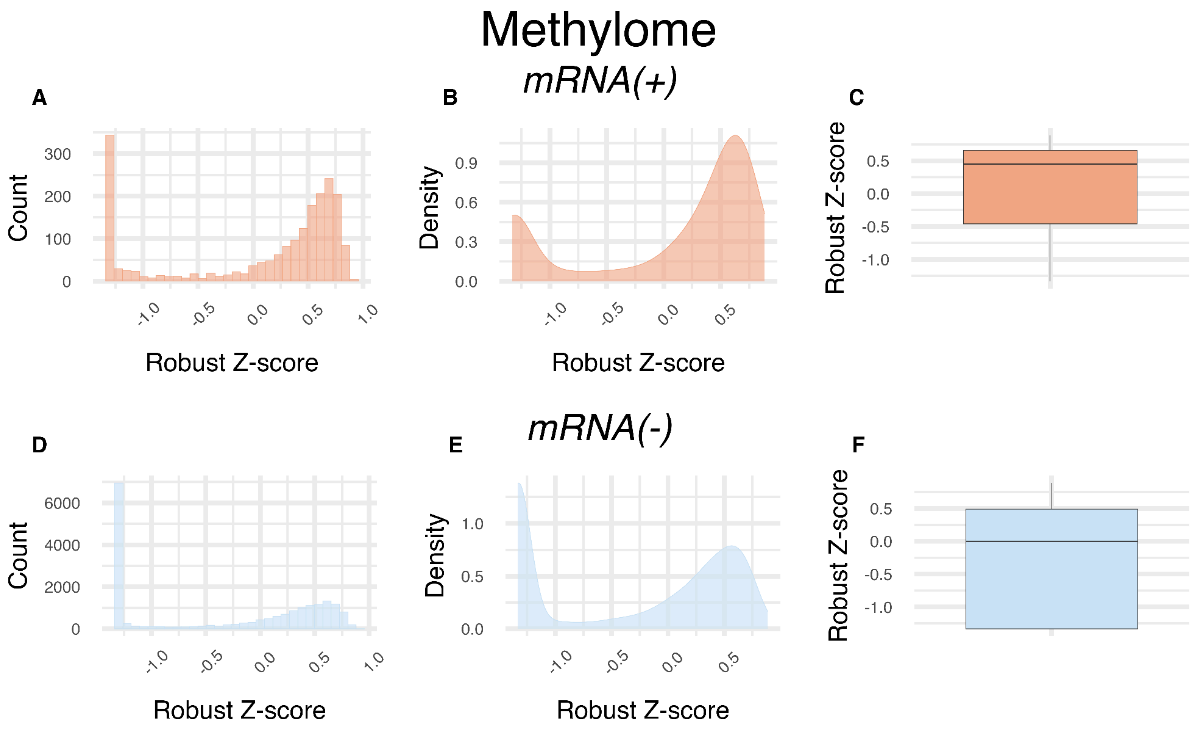


**Supplementary Figure S7. mRNA methylome robust z-scores distributions in log scale.** A-C) Histogram, Density and Boxplot for mRNA(+), D-F) Histogram, Density and Boxplot for mRNA(-) robust z-scores computed from the average proportion of methylated CpG sites across samples.

### Selection of most relevant epigenetic features

A Spearman correlation matrix was generated to assess correlations among epigenetic features and identify the most informative and uncorrelated variables for subsequent analysis. Kolmogorov-Smirnov statistics were calculated to quantify distributional differences between positive and negative control sequences, enabling identification of features with the greatest discriminatory power. These plots were created using the R functions ‘rcorr’ from Hmisc v.5.2-3 package, ‘heatmaply’ from Heatmaply v1.5.0 package, ‘p.adjust’ and ‘ks.test’ from Stats v.4.4.1 package

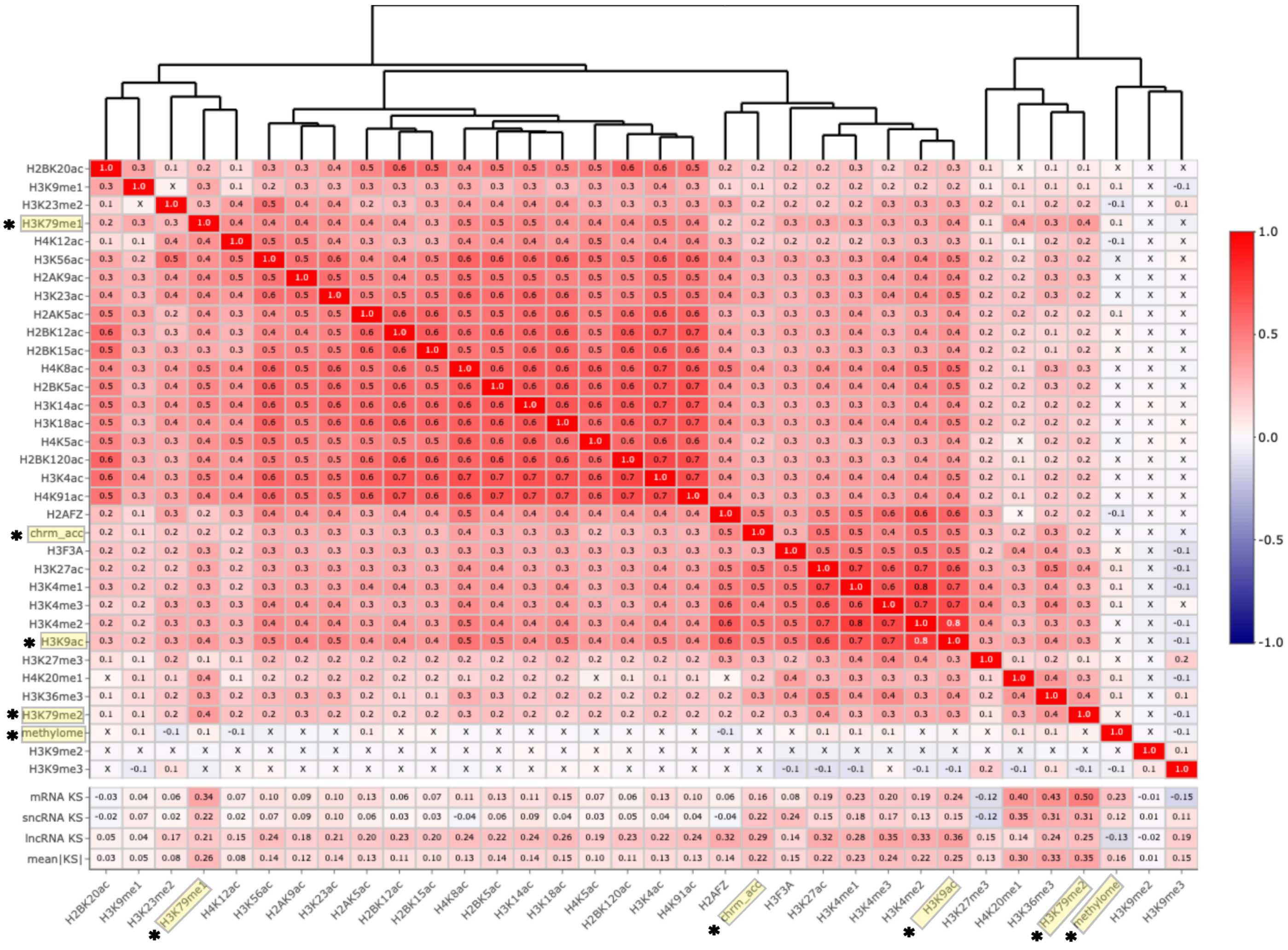


**Supplementary Figure S8. Correlation matrix of epigenetic features.** Spearman correlation of histone and DNA modifications for mRNA, sncRNA and lncRNA sequences. Positively correlated features are displayed in red and negatively correlated features in blue. Spearman’s Rho was used to determine the significance of each correlation, which were corrected for multiple testing using a Benjamini-Hochberg (BH) correction. Non-significant correlations with a p-value > 0.05 have been marked with a cross. At the bottom of the plot, KS statistics are included for each gene type as well as a mean |KS| for overall comparison.

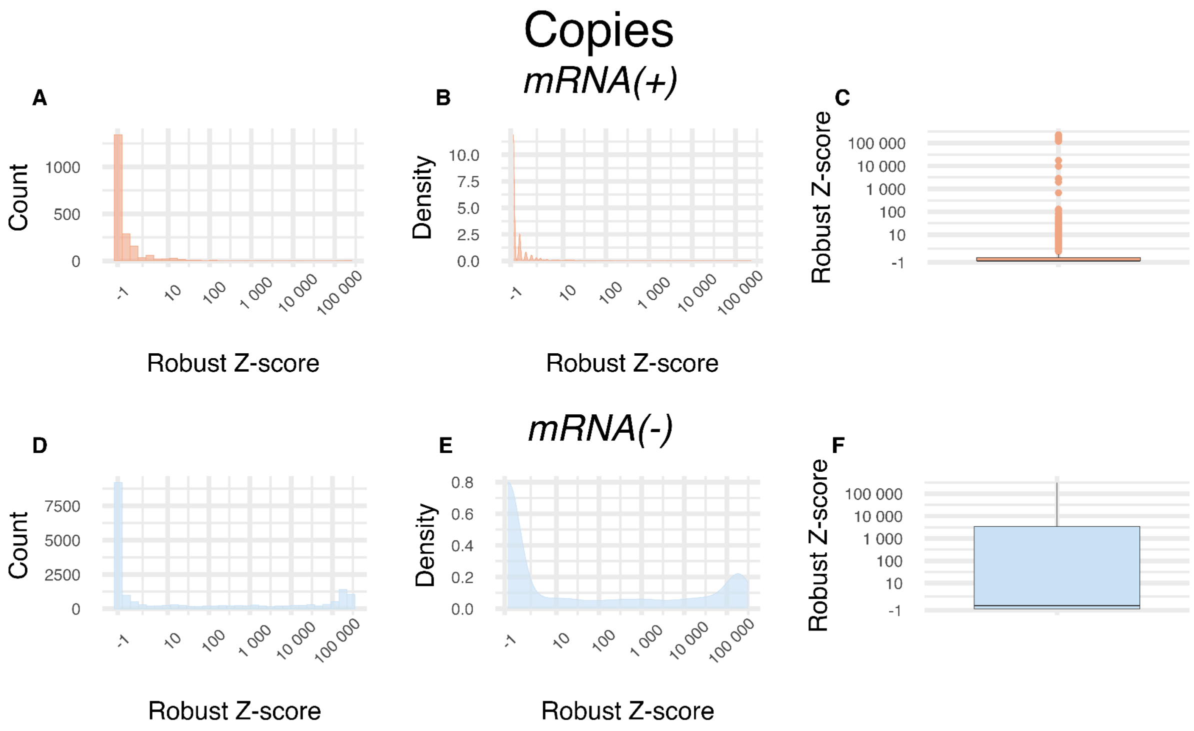


**Supplementary Figure S9. mRNA copies robust z-scores distributions in log scale.** A-C) Histogram, Density and Boxplot for mRNA(+), D-F) Histogram, Density and Boxplot for mRNA(-) of z-scores computed from the number of genomic copies of each region.

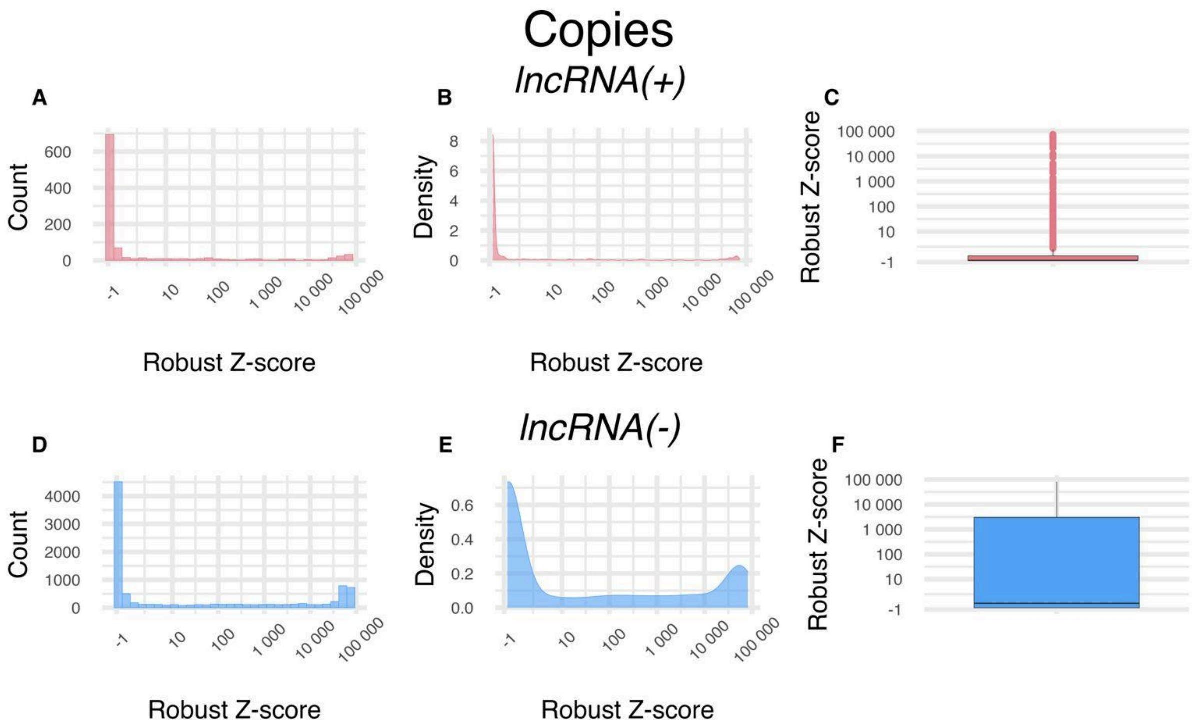


**Supplementary Figure S10. lncRNA copies robust z-scores distributions in log scale.** A-C) Histogram, Density and Boxplot for lncRNA(+), D-F) Histogram, Density and Boxplot for lncRNA(-) of z-scores computed from the number of genomic copies of each region.

### Effect on distance

In order to demonstrate that there was no effect on distance to gene for the negative control sequences, Spearman correlation coefficients were calculated between robust z-scores using the R functions rcorr from Hmisc v. 5.2-3 package. Then to create the heatscatter plots the function heatscatter from LSD v. 4.1-0 package was used, and to compute the trend line the loess function from stats v. 4.4.1 package was used.

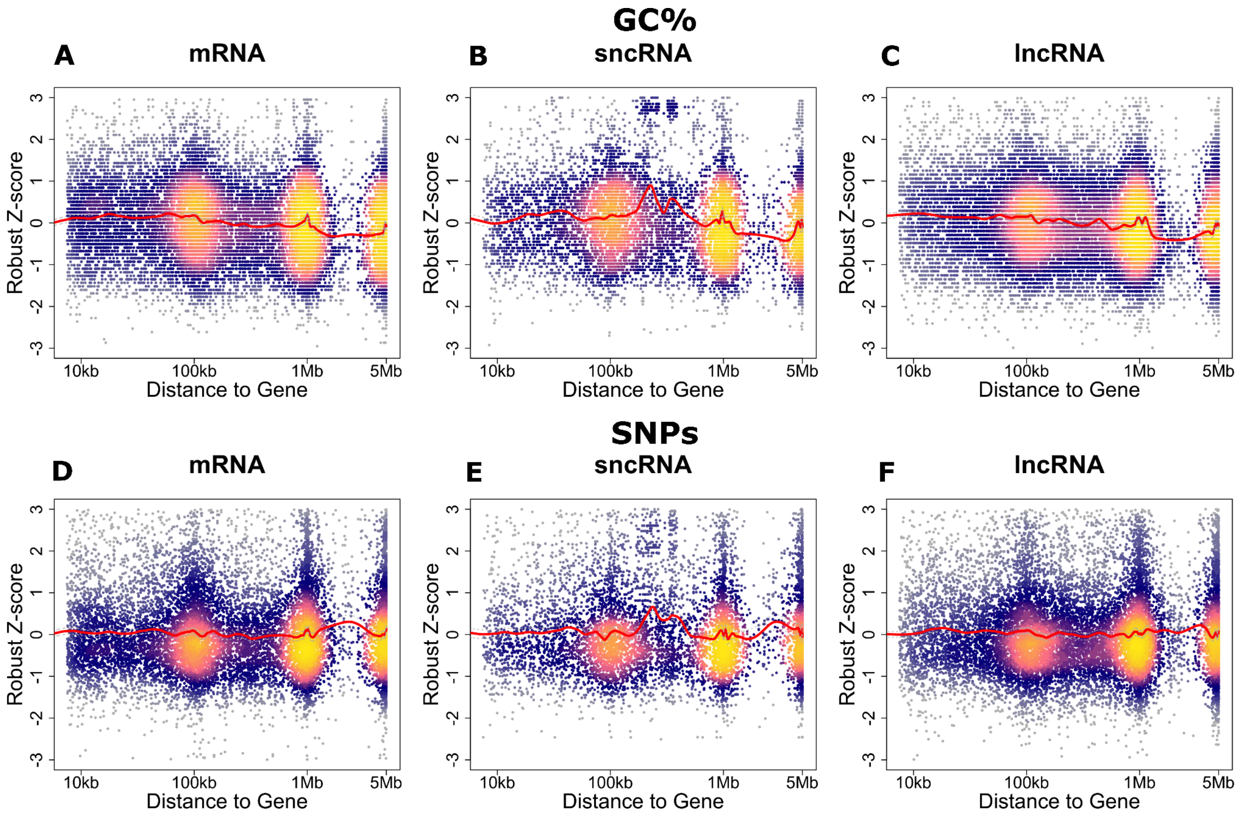


**Supplementary Figure S11. Heatscatter for the effect on distance.** A-C) GC% for each gene type, D-F) SNPs for each gene type. Robust Z-scores are displayed on the y-axis as a function of the distance to the nearest gene in the x-axis. Each point represents a sequence. There is no clear linear relationship between distance to the gene and GC content variation or SNPs.

We generated an Upset plot to determine which genome annotation databases should be used to select negative control sequences that corresponded to intergenic unannotated regions of the genome. The R function ‘upset’ from the UpSetR v1.4.0 package was used. The used script (“UpsetPlot.R”) is included in the project repository. The databases that contained the highest number of annotations (GencodeV45) and unique annotations (RNACentral) were selected, as they encapsulated the most annotations.

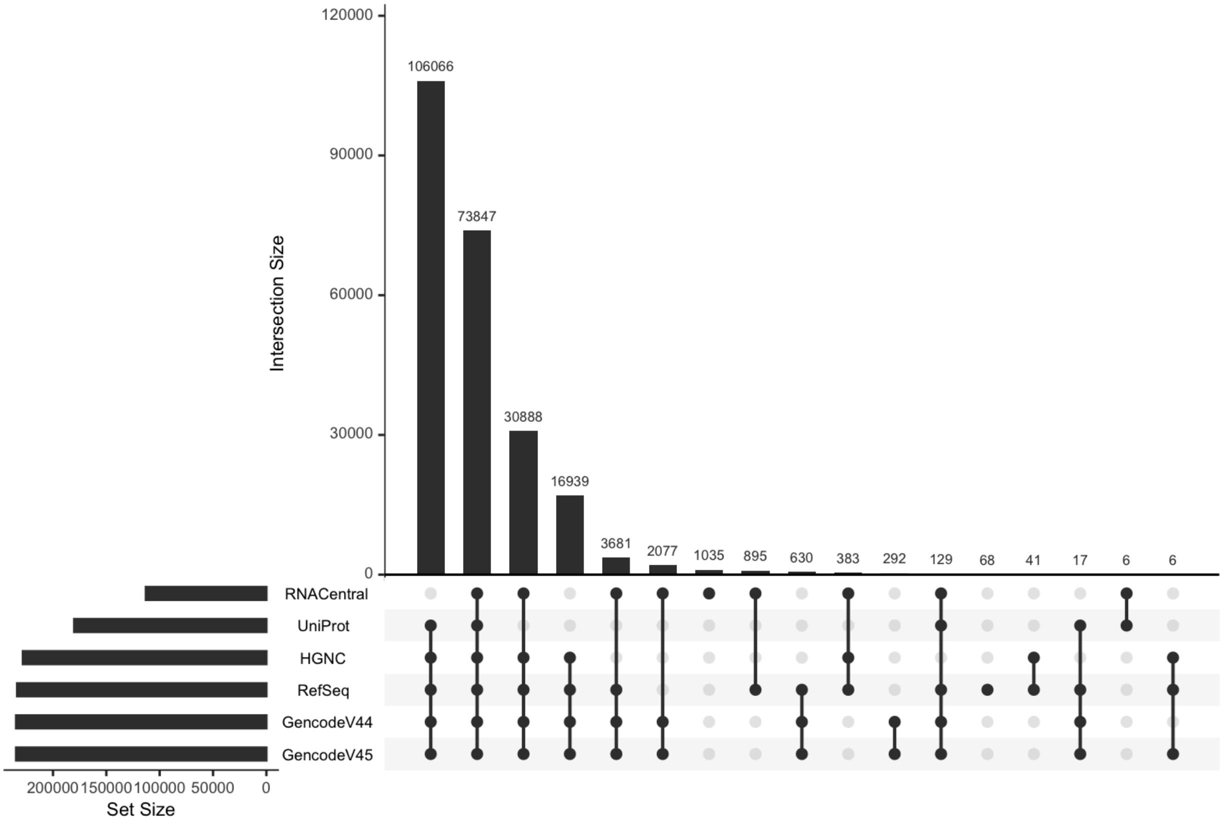


**Supplementary Figure S12. Upset plot for different human genome databases (build GRCh38) corresponding to annotations from chromosome 22.** This figure illustrates the overlap of gene annotations from six prominent databases: RNACentral, UniProt, HGNC, RefSeq, Gencode V44, and Gencode V45. The horizontal bars represent the total number of gene annotations in each database. The vertical bars represent the number of gene annotations that are common (or unique) across the selected combinations of databases, indicated by the connected dots below. The connected dots indicate the specific databases involved in each intersection.

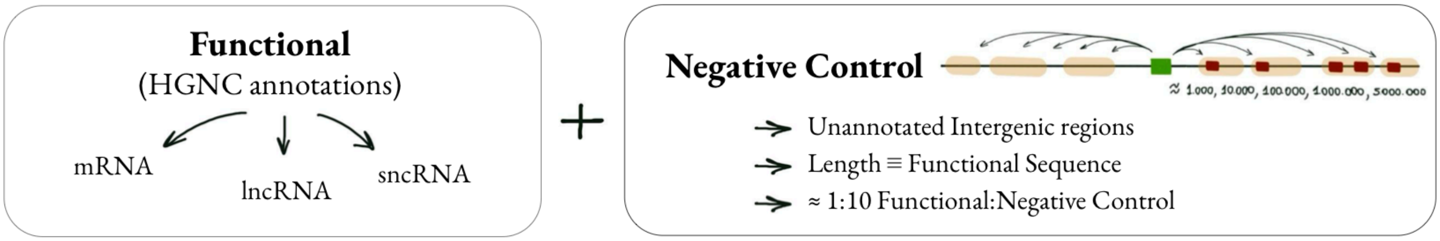


**Supplementary Figure S13. Workflow for generating control datasets.** Functional sequences were selected from HGNC-linked mRNA, lncRNA, and sncRNA annotations. For each functional sequence, length-matched negative controls were sampled from unannotated intergenic regions of the GRCh38 genome at increasing distances from the corresponding gene (approximately 1, 10, 100, 1000, and 5000 kb upstream and downstream). Candidate controls were filtered to remove overlaps with known annotations, generating a final set of negative controls at an approximate 1:10 functional-to-control ratio.

### Result summaries/Data overviews

To provide an overall multivariate view of the data, we performed a principal component analysis (PCA) on the robust z-scores of the most informative genomic features. Unlike the per-feature analyses presented in the main text, PCA summarises how features act in combination, allowing us to assess whether functional and non-genic sequences separate in reduced-dimensional space and to identify which variables contribute most strongly to that separation. These plots therefore serve as a complementary overview of the structure, redundancy, and discriminatory power of the selected feature sets across mRNA, sncRNA, and lncRNA datasets.

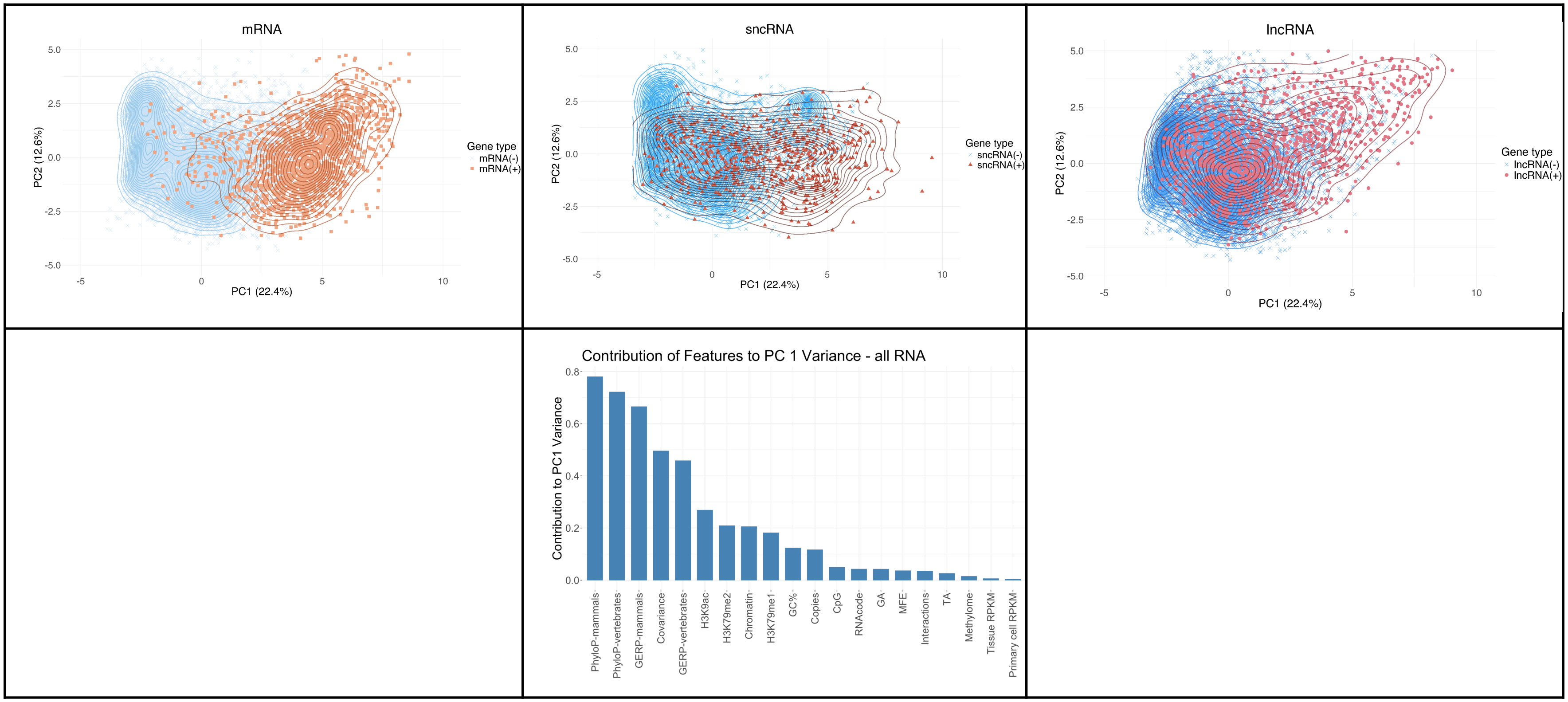
mRNA
5.0
2.5
0.0
-2.5
-5.0
PC2 (12.6%)
-5
0
5
10
PC1 (22.4%)
Gene type
mRNA(-)
mRNA(+)
sncRNA
5.0
2.5
0.0
-2.5
-5.0
PC2 (12.6%)
-5
0
5
10
PC1 (22.4%)
Gene type
sncRNA(-)
sncRNA(+)
lncRNA
5.0
2.5
0.0
-2.5
-5.0
PC2 (12.6%)
-5
0
5
10
PC1 (22.4%)
Gene type
lncRNA(-)
lncRNA(+)
Contribution of Features to PC 1 Variance - all RNA
0.8
0.6
0.4
0.2
0.0
Contribution to PC1 Variance
PhyloP-mammals
PhyloP-vertebrates
GERP-mammals
Covariance
GERP-vertebrates
H3K9ac
H3K79me2
Chromatin
H3K79me1
GC%
Copies
CpG
RNAcode
GA
MFE
Interactions
TA
Methylome
Tissue RPKM
Primary cell RPKM

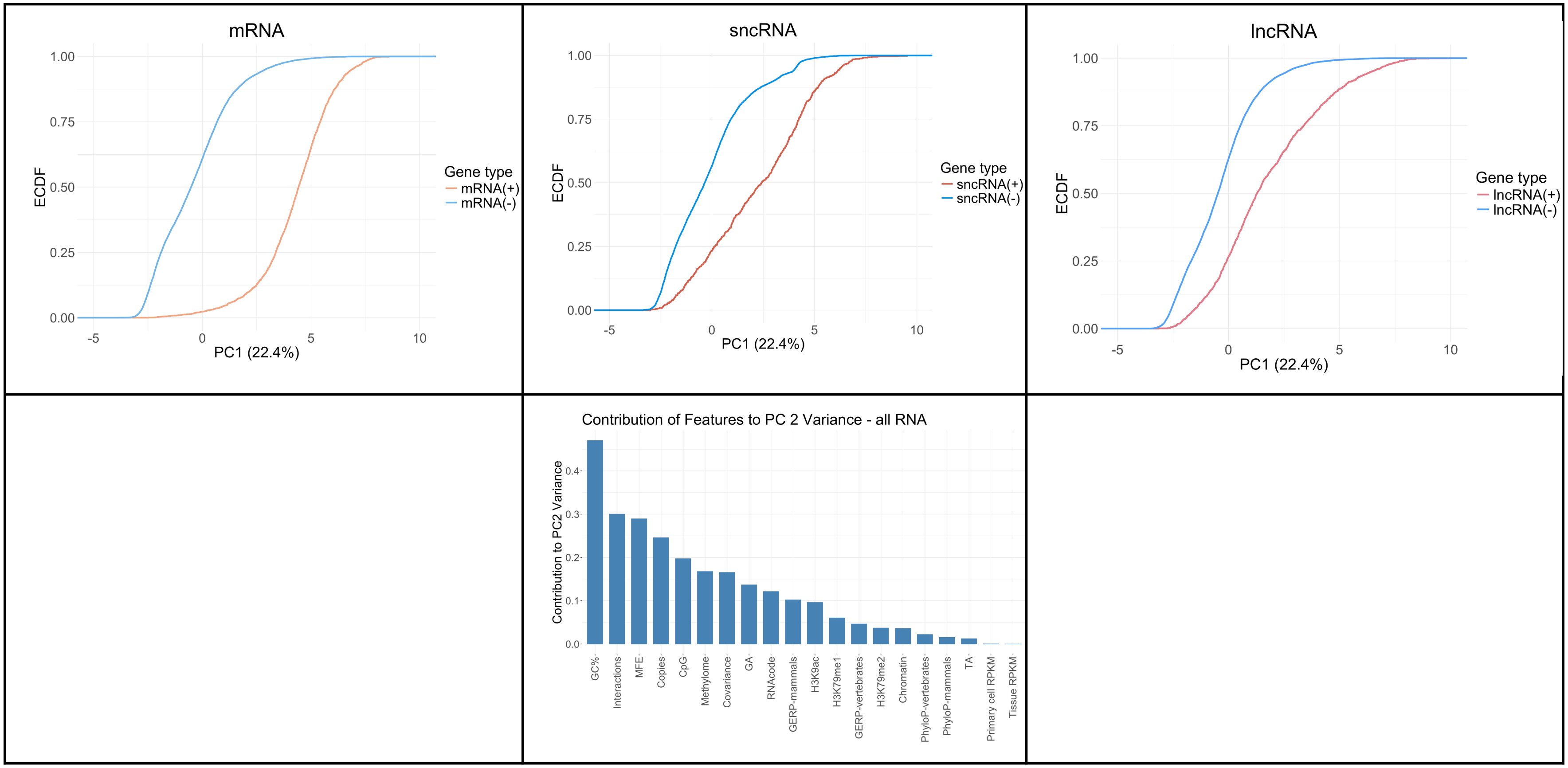
mRNA
ECDF
1.00
0.75
0.50
0.25
0.00
-5
0
5
10
PC1 (22.4%)
Gene type
mRNA(+)
mRNA(-)
sncRNA
Gene type
sncRNA(+)
sncRNA(-)
lncRNA
Gene type
lncRNA(+)
lncRNA(-)
Contribution of Features to PC 2 Variance - all RNA
Contribution to PC2 Variance
0.4
0.3
0.2
0.1
0.0
GC%
Interactions
MFE
Copies
CpG
Methylome
Covariance
GA
RNAcode
GERP-mammals
H3K9ac
H3K79me1
GERP-vertebrates
H3K79me2
Chromatin
PhyloP-vertebrates
PhyloP-mammals
TA
Primary cell RPKM
Tissue RPKM

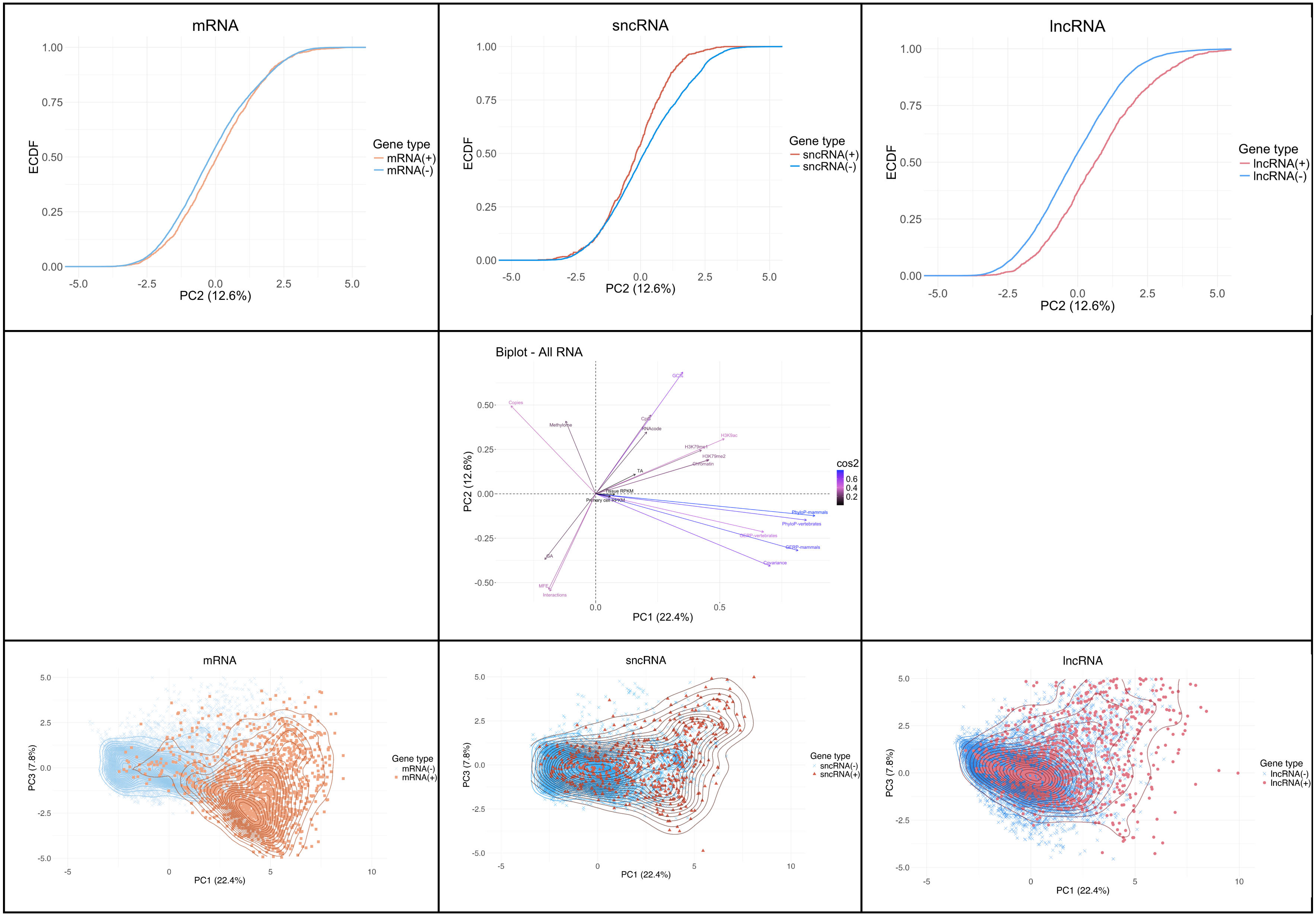
mRNA
ECDF
PC2 (12.6%)
Gene type
mRNA(+)
mRNA(-)
sncRNA
sncRNA(+)
sncRNA(-)
lncRNA
lncRNA(+)
lncRNA(-)
Biplot - All RNA
PC2 (12.6%)
PC1 (22.4%)
cos2
Copies
Methylome
GC%
CpG
RNAcode
H3K9ac
H3K79me1
H3K79me2
Chromatin
TA
Tissue RPKM
Primary cell RPKM
PhyloP-mammals
PhyloP-vertebrates
GERP-vertebrates
GERP-mammals
Covariance
GA
MFE
Interactions
PC3 (7.8%)
PC1 (22.4%)

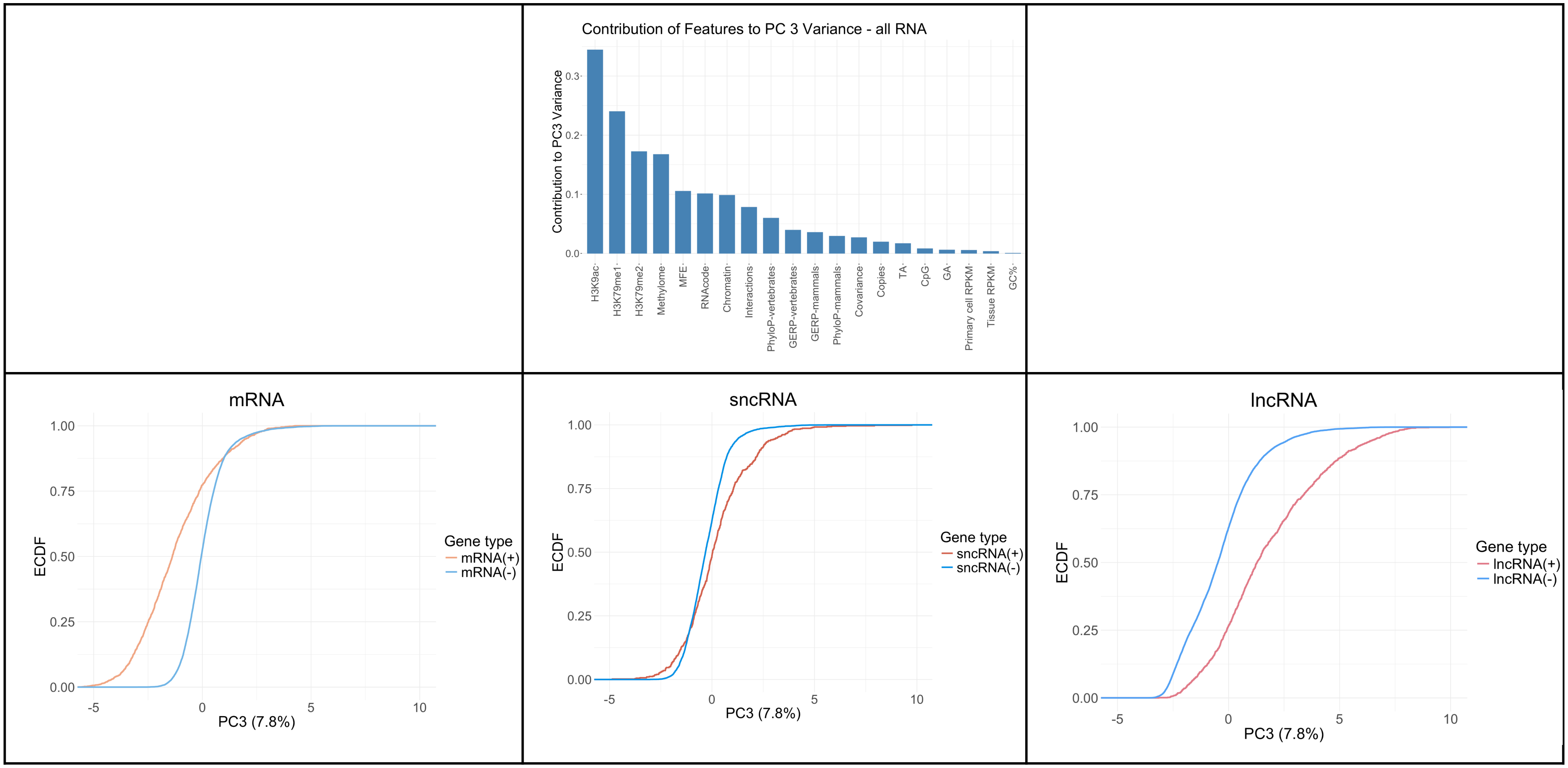

Contribution of Features to PC 3 Variance - all RNA
Contribution to PC3 Variance
0.0
0.1
0.2
0.3
H3K9ac
H3K79me1
H3K79me2
Methylome
MFE
RNAcode
Chromatin
Interactions
PhyloP-vertebrates
GERP-vertebrates
GERP-mammals
PhyloP-mammals
Covariance
Copies
TA
CpG
GA
Primary cell RPKM
Tissue RPKM
GC%
mRNA
ECDF
PC3 (7.8%)
Gene type
mRNA(+)
mRNA(-)
sncRNA
sncRNA(+)
sncRNA(-)
lncRNA
lncRNA(+)
lncRNA(-)
0.00
0.25
0.50
0.75
1.00
-5
0
5
10

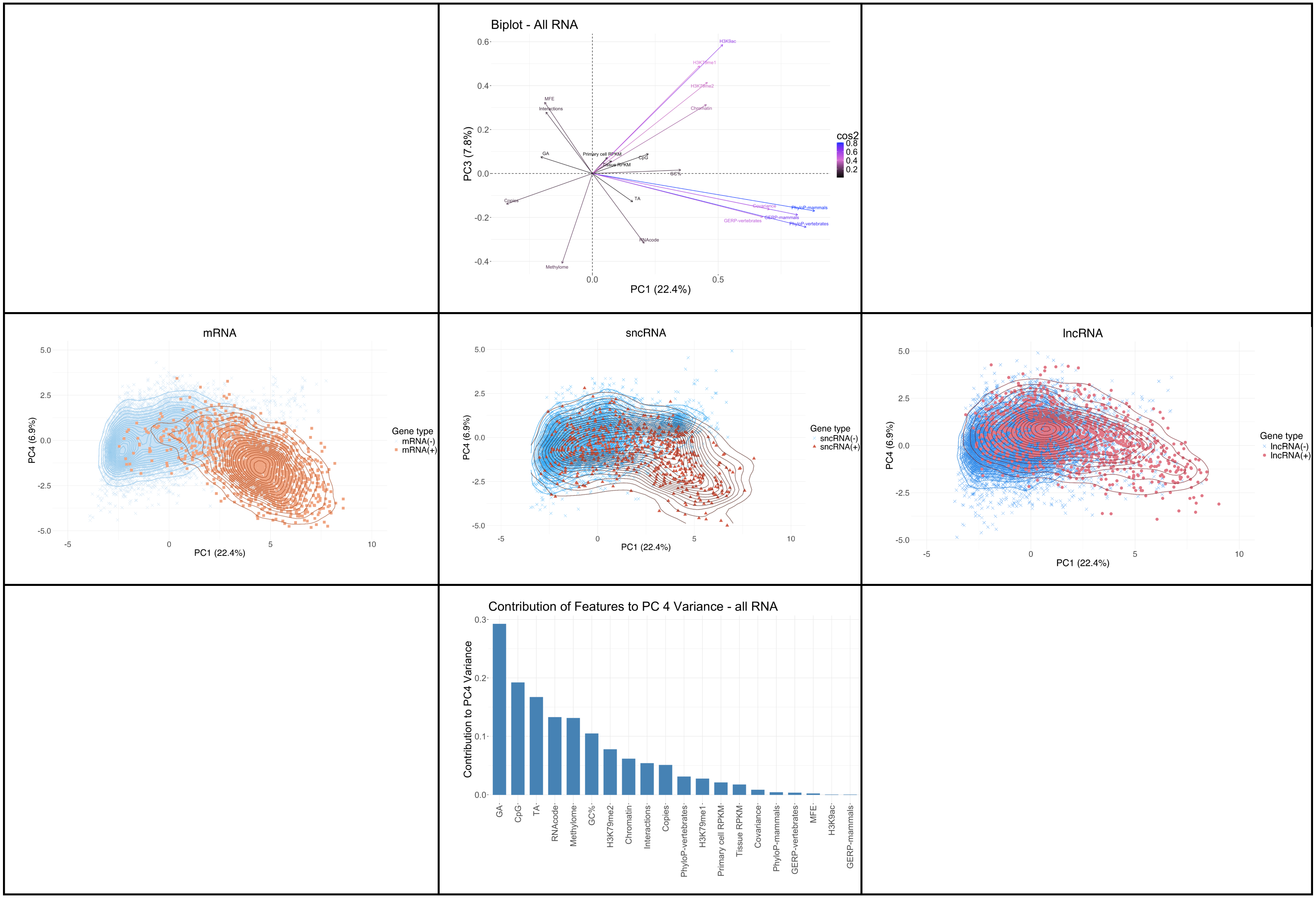
Biplot - All RNA
PC3 (7.8%)
PC1 (22.4%)
cos2
0.8
0.6
0.4
0.2
H3K9ac
H3K79me1
H3K79me2
Chromatin
MFE
Interactions
Primary cell RPKM
GA
CpG
Tissue RPKM
TA
Copies
Covariance
PhyloP-mammals
GERP-mammals
GERP-vertebrates
PhyloP-vertebrates
RNAcode
Methylome
mRNA
sncRNA
lncRNA
PC4 (6.9%)
Gene type
mRNA(-)
mRNA(+)
sncRNA(-)
sncRNA(+)
lncRNA(-)
lncRNA(+)
Contribution of Features to PC 4 Variance - all RNA
Contribution to PC4 Variance
GA
CpG
TA
RNAcode
Methylome
GC%
H3K79me2
Chromatin
Interactions
Copies
PhyloP-vertebrates
H3K79me1
Primary cell RPKM
Tissue RPKM
Covariance
PhyloP-mammals
GERP-vertebrates
MFE
H3K9ac
GERP-mammals

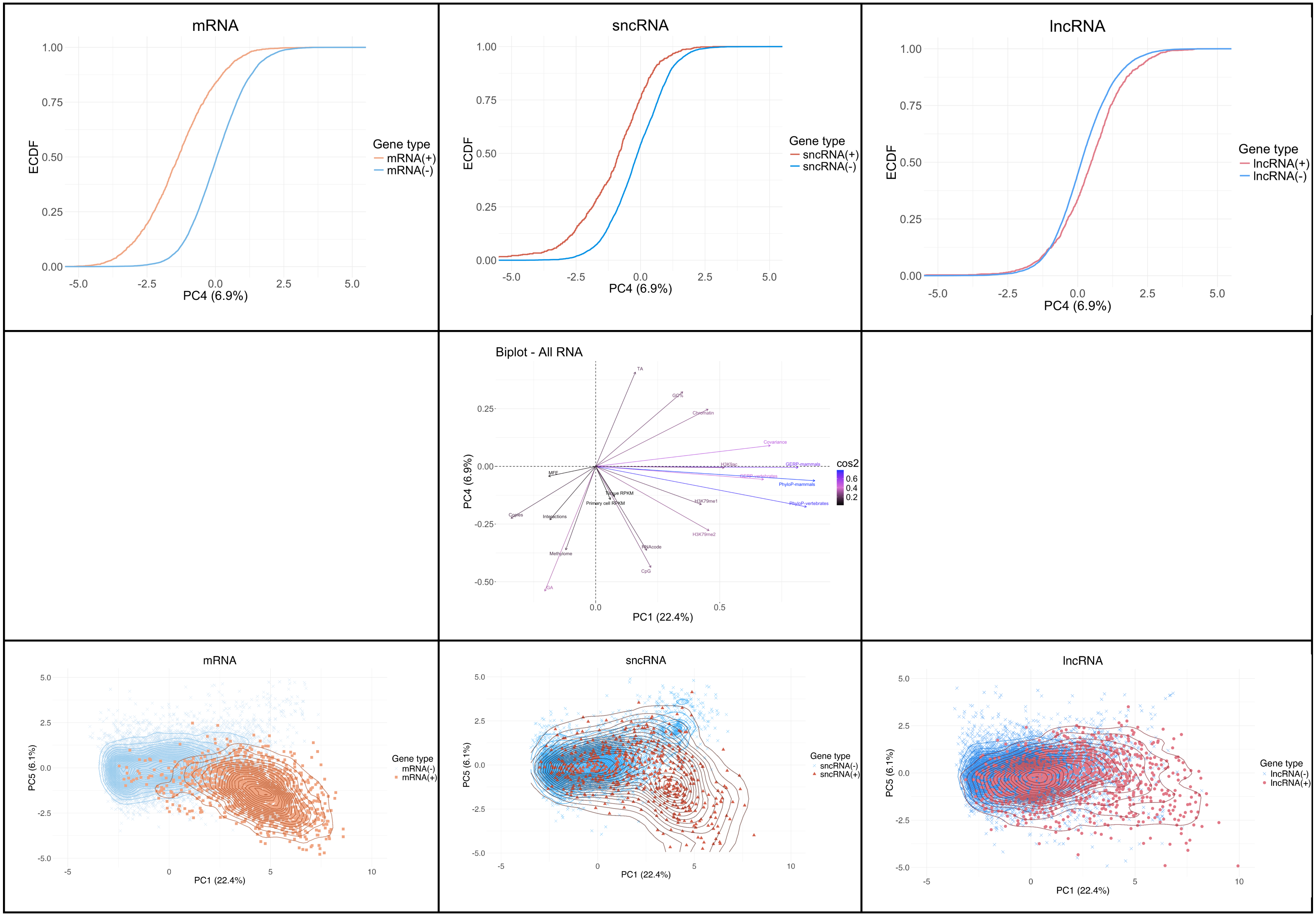
mRNA
ECDF
PC4 (6.9%)
Gene type
mRNA(+)
mRNA(-)
sncRNA
sncRNA(+)
sncRNA(-)
lncRNA
lncRNA(+)
lncRNA(-)
Biplot - All RNA
PC4 (6.9%)
PC1 (22.4%)
cos2
TA
GC%
Chromatin
Covariance
H3K9ac
GERP-mammals
GERP-vertebrates
PhyloP-mammals
PhyloP-vertebrates
H3K79me1
H3K79me2
RNAcode
CpG
MFE
Tissue RPKM
Primary cell RPKM
Copies
Interactions
Methylome
GA
PC5 (6.1%)
PC1 (22.4%)

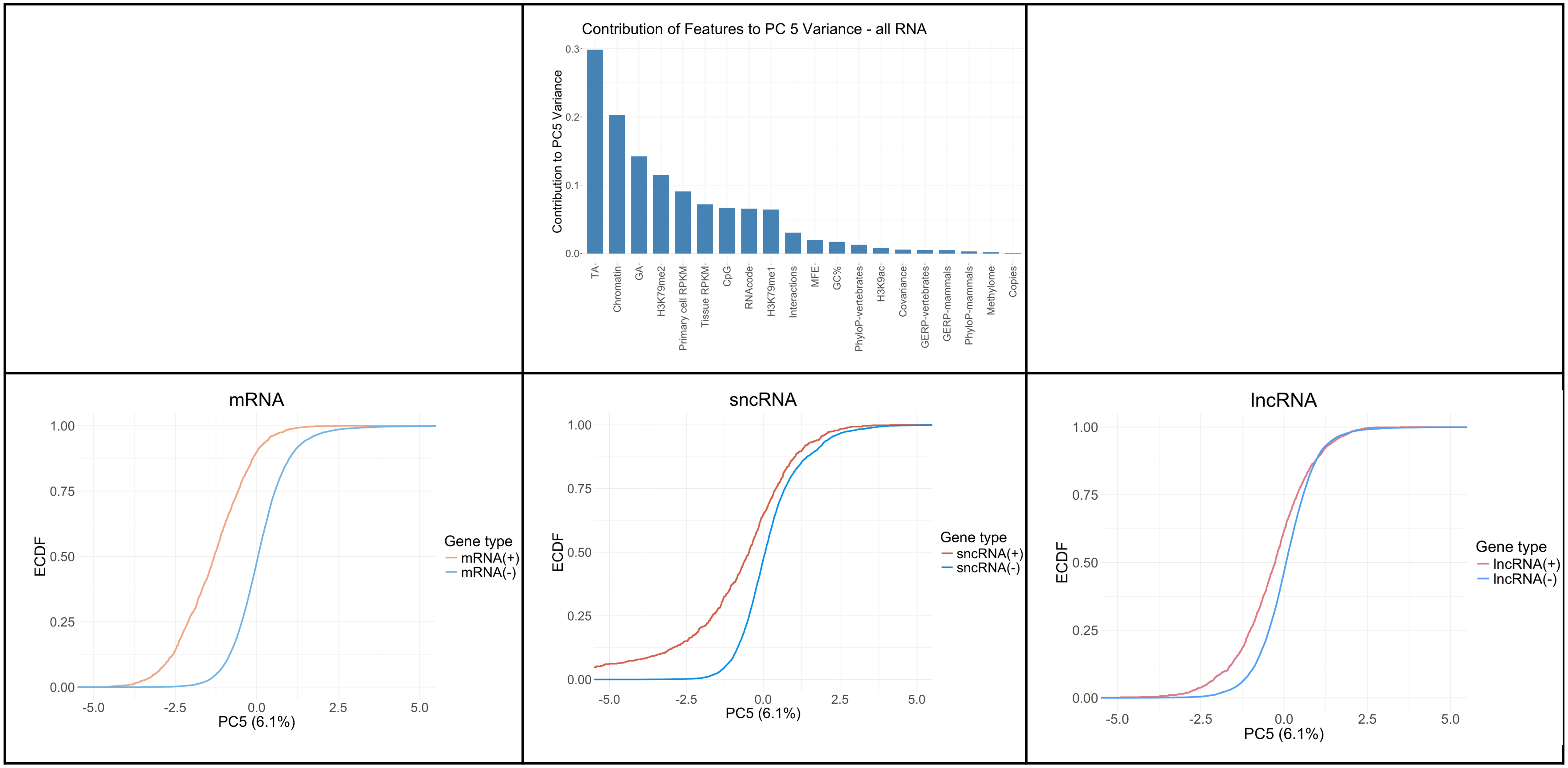

Contribution of Features to PC 5 Variance - all RNA
Contribution to PC5 Variance
0.3
0.2
0.1
0.0
TA
Chromatin
GA
H3K79me2
Primary cell RPKM
Tissue RPKM
CpG
RNAcode
H3K79me1
Interactions
MFE
GC%
PhyloP-vertebrates
H3K9ac
Covariance
GERP-vertebrates
GERP-mammals
PhyloP-mammals
Methylome
Copies
mRNA
ECDF
1.00
0.75
0.50
0.25
0.00
-5.0
-2.5
0.0
2.5
5.0
PC5 (6.1%)
Gene type
mRNA(+)
mRNA(-)
sncRNA
sncRNA(+)
sncRNA(-)
lncRNA
lncRNA(+)
lncRNA(-)

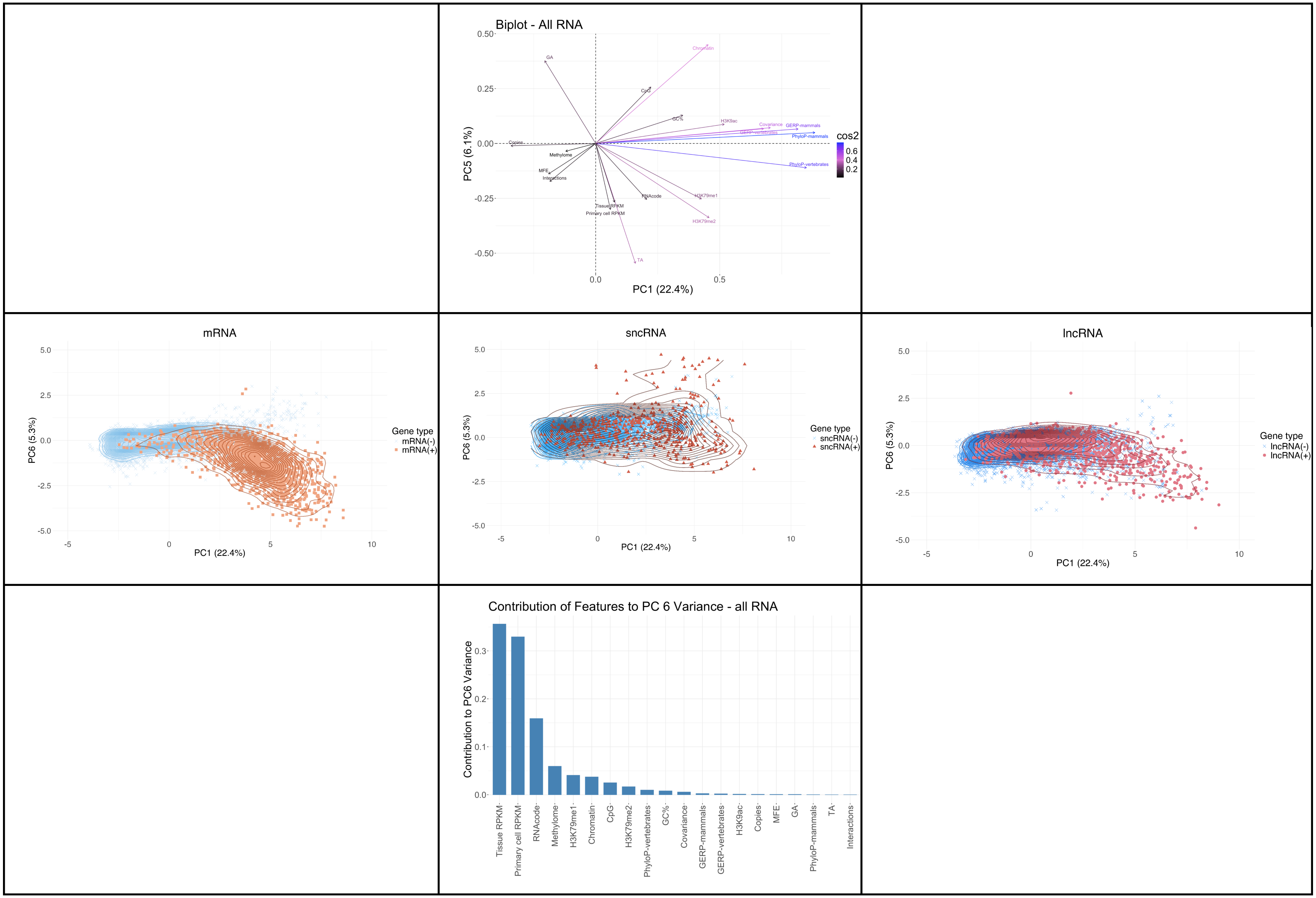
Biplot - All RNA
PC5 (6.1%)
PC1 (22.4%)
cos2
0.6
0.4
0.2
GA
Chromatin
CpG
GC%
H3K9ac
Covariance
GERP-mammals
GERP-vertebrates
PhyloP-mammals
PhyloP-vertebrates
Copies
Methylome
MFE
Interactions
RNAcode
H3K79me1
H3K79me2
Tissue RPKM
Primary cell RPKM
TA
mRNA
sncRNA
lncRNA
PC6 (5.3%)
PC1 (22.4%)
Gene type
mRNA(-)
mRNA(+)
sncRNA(-)
sncRNA(+)
lncRNA(-)
lncRNA(+)
Contribution of Features to PC 6 Variance - all RNA
Contribution to PC6 Variance
Tissue RPKM
Primary cell RPKM
RNAcode
Methylome
H3K79me1
Chromatin
CpG
H3K79me2
PhyloP-vertebrates
GC%
Covariance
GERP-mammals
GERP-vertebrates
H3K9ac
Copies
MFE
GA
PhyloP-mammals
TA
Interactions

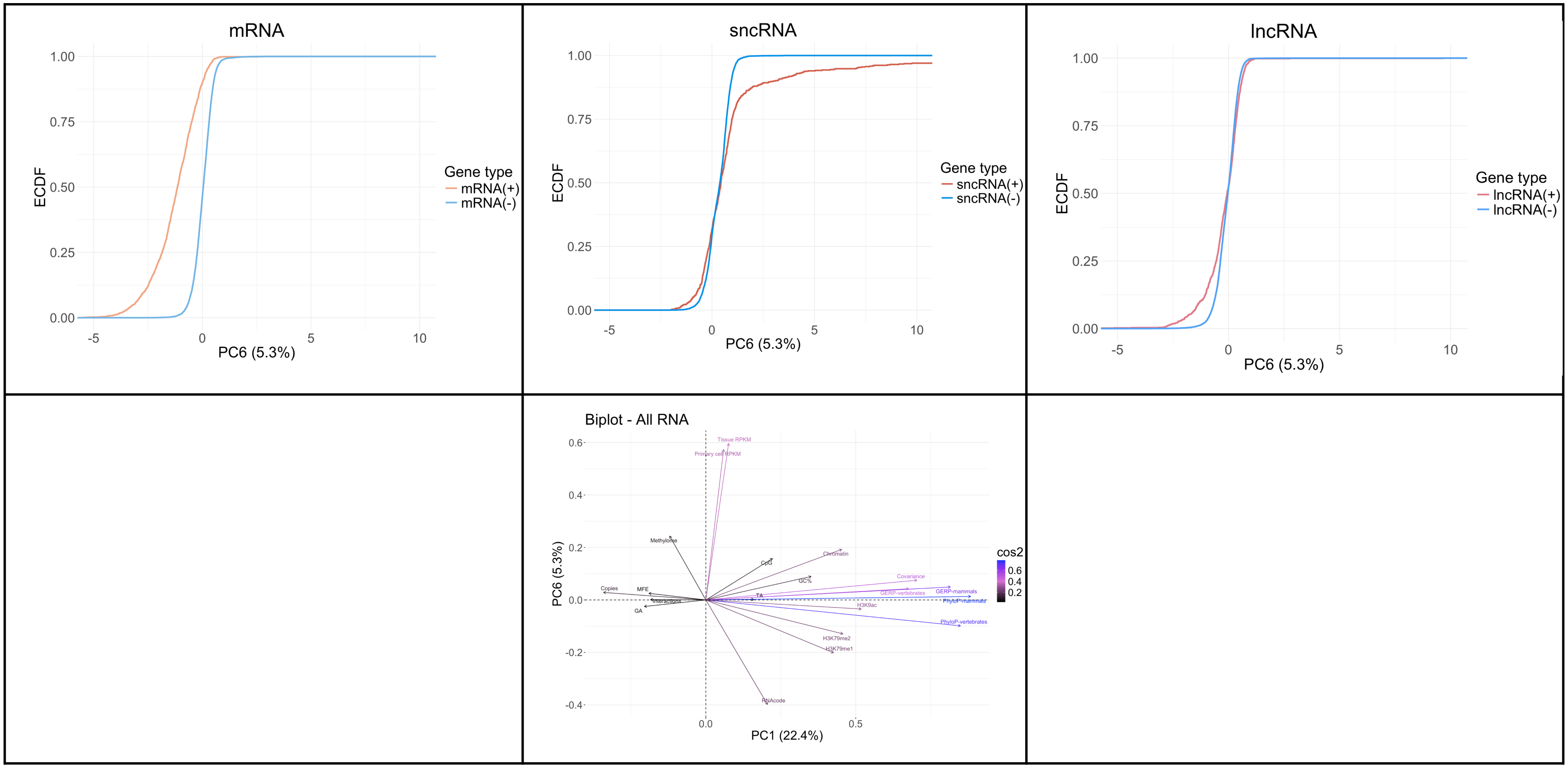


**Fig S14. Principal Component Analysis (PCA).** The plots include scatter and contour plots, screeplot, univariate plot, and biplot for PC1-PC6. The scatter and contour plots show the clustering of positive and negative sequences for each gene type between PC1 vs PC2-PC6. Screeplot indicates the contribution of each genomic feature to its corresponding PC. The univariate plot illustrates the empirical cumulative distribution function (ECDF) for each PC divided by gene type. Biplots show feature loadings in PC1 vs PC2-PC6 space, vector arrows indicate the direction and magnitude of feature contributions to each principal component.
